# Molecular insight into the *Drosophila* piRNA pathway network through a combination of systematic protein interaction screening and structural prediction

**DOI:** 10.1101/2024.05.31.596839

**Authors:** Harpreet Kaur Salgania, Jutta Metz, Eric Lingren, Christian Bleischwitz, David Hauser, Katalin Oliveras Máté, Daniel Bollack, Felix Lahr, Asen Garbelyanski, Mandy Jeske

## Abstract

piRNA-bound PIWI proteins mediate the silencing of transposons at both the transcriptional and post-transcriptional levels, processes that are critical for genome integrity and fertility in animals. While numerous additional proteins are known to be essential for piRNA biogenesis and function in *Drosophila* and other animals, their molecular and mechanistic functions have remained largely unknown. To improve our molecular understanding of the *Drosophila* piRNA pathway, we used a cell culture-based protein-protein interaction assay called ReLo to perform a systematic pairwise interaction screen involving 22 factors operating in the cytoplasm, including PIWI proteins, Tudor domain-containing proteins (TDRDs), RNA helicases, and mitochondrial surface proteins. Through additional ReLo interaction testing and structural modeling using AlphaFold-Multimer, we have characterized six protein complexes at the molecular and structural levels. We believe that the results of this screen and our methodological approach are likely to guide future research into the molecular mechanisms underlying piRNA biogenesis and function.

## INTRODUCTION

Piwi-interacting RNAs are a class of short non-coding RNAs expressed predominantly in the gonads of animals. In complex with PIWI proteins, piRNAs recognize their targets based on sequence complementarity and mediate the silencing of transposons and other genes at both transcriptional and post-transcriptional levels, thereby maintaining genome integrity and fertility in animals (Haase, 2016; Czech & Hannon, 2016; Huang et al, 2017; Hirakata & Siomi, 2016; Yamashiro & Siomi, 2018).

The piRNA pathway has been extensively studied in the *Drosophila* ovary (Handler et al, 2013; Czech et al, 2013; Olivieri et al, 2010; Huang et al, 2017), where piRNAs are expressed in both somatic and germline tissues. In both tissues, piRNAs are encoded by piRNA clusters, which are intergenic DNA segments rich in fragmented transposon sequences (Vagin et al, 2006; Brennecke et al, 2007; Yamanaka et al, 2014). After synthesis in the nucleus, piRNA precursors are exported to the cytoplasm, where they are loaded onto PIWI proteins and processed into mature piRNAs.

*Drosophila* ovaries express three PIWI proteins, each with a specialized function. The germline-specific piRNA-bound PIWI proteins Aubergine (Aub) and Argonaute 3 (Ago3) recognize and catalyze the cleavage of target mRNAs in the cytoplasm as part of the piRNA amplification loop, also known as the ping-pong cycle (Brennecke et al, 2007; Gunawardane et al, 2007). This cycle involves the reciprocal cleavage of sense and antisense piRNA precursors, thereby amplifying piRNA production. In contrast, piRNA-bound Piwi, which lacks cleavage activity, induces transcriptional repression of targeted transposon genes in the nucleus (Darricarrére et al, 2013; Sienski et al, 2012; Le Thomas et al, 2014a, 2014b). Piwi-bound piRNAs are generated through phasing, a process involving the mitochondrially localized endonuclease Zucchini (Zuc), which cleaves piRNA precursors into phased piRNAs (Ipsaro et al, 2012; Haase et al, 2010; Nishimasu et al, 2012; Han et al, 2015; Mohn et al, 2015). Phasing generates a series of piRNAs from a single precursor transcript, increasing the diversity of piRNA sequences and expanding the repertoire of transposon targets.

In addition to the PIWI proteins, *Drosophila* and other animals express a variety of other critical components of the piRNA pathway, including RNA helicases and Tudor domain-containing proteins (TDRDs) (Olivieri et al, 2010; Handler et al, 2013; Czech et al, 2013; Ozata et al, 2019). Similar to the loss of PIWI proteins, disruption of the *in vivo* function of the additional piRNA factors results in defects in piRNA production and failure to effectively repress transposons in the gonads, ultimately leading to infertility of the organism. The roles of many of these additional piRNA factors have been assigned to specific steps within the pathway, such as piRNA precursor recognition, piRNA amplification, or phasing. However, for most of these proteins the molecular mechanisms underlying their function remain largely unknown.

Most ovarian cytoplasmic piRNA factors assemble into electron-dense, granular cytoplasmic structures. Ovarian follicle cells, which are of somatic origin, contain Yb bodies that accumulate factors required for piRNA precursor recognition (Szakmary et al, 2009; Pandey et al, 2017; Murota et al, 2014; Ishizu et al, 2015; Saito et al, 2009; Homolka et al, 2015; Rogers et al, 2017). In contrast, nurse cells, which are of germline origin, contain perinuclear nuage that accumulates factors critical for the ping-pong cycle (Lim & Kai, 2007). The granular localization of many piRNA factors suggests their structural complexity and poor accessibility for study by biochemical methods.

Nevertheless, experimental structures have been determined for several individual piRNA factors or protein domains that function in the cytoplasm or at the mitochondrial surface. These include the PIWI protein cores in the apo form, bound to piRNA, or bound to piRNA and a target sequence (Matsumoto et al, 2016; Yamaguchi et al, 2020; Anzelon et al, 2021), the core of the Vasa DEAD-box RNA helicase bound to a non-hydrolyzable ATP analog and an RNA oligo (Sengoku et al, 2006; Xiol et al, 2014), the nuclease domains of Maelstrom (Matsumoto et al, 2015; Chen et al, 2015) and Zucchini (Nishimasu et al, 2012; Voigt et al, 2012; Ipsaro et al, 2012), and several isolated eTudor domains from different TDRDs (Liu et al, 2010a; Mathioudakis et al, 2012; Ren et al, 2014; Zhang et al, 2018, 2017).

In contrast, experimental structural information on cytoplasmic protein complexes is sparse and can essentially be summarized by two characterized interaction modules. One module is formed between the C-terminal RecA-like domain of the ATP-dependent RNA helicase Vasa and an eLOTUS domain (Jeske et al, 2017) (**Supplementary Figure 1A**), which, in *Drosophila,* is present in the germline inducer Oskar and in the piRNA factors Tejas (TDRD5) and Tapas (TDRD7) (Anantharaman et al, 2010; Callebaut & Mornon, 2010; Kubíková et al, 2020). eLOTUS domains bind and stimulate the ATPase activity of Vasa *in vitro* (Jeske et al, 2017), but the biological function of this stimulation remains unclear. The other module includes eTudor domains, which were previously described to bind to sequence motifs containing symmetrically dimethylated arginine residues (sDMA) found in the arginine/glycine-rich, unstructured N-terminal extensions of PIWI proteins (Kirino et al, 2009; Liu et al, 2010a; Zhang et al, 2018; Huang et al, 2021; Mathioudakis et al, 2012; Liu et al, 2010b). Several crystal structures of eTudor domains in complex with a short PIWI peptide revealed the modes of sDMA-dependent (and sDMA-independent) interactions (Liu et al, 2010a; Zhang et al, 2018; Huang et al, 2021; Zhang et al, 2017; Chen et al, 2020; Liu et al, 2010b) (**Supplementary Figure 1B**). To recognize the positively charged arginine-rich ligands, eTudor domains use a negatively charged cleft, formed by the canonical Tudor domain part and an SN-like domain part, in which the sDMA is accommodated by an aromatic cage composed of four aromatic residues emanating from the Tudor domain part (Gan et al, 2019). In *Drosophila*, sDMA-dependent interactions have been reported for the eTudor domains present in the TDRDs Tudor, Papi, and Krimper (Liu et al, 2010a; Zhang et al, 2018; Huang et al, 2021). However, the eTudor domains of many other piRNA factors lack an intact aromatic cage (**Supplementary Figure 1C**). Although there is an example of an eTudor domain lacking an intact aromatic cage binding to a peptide containing unmethylated arginine residues (Huang et al, 2021) (**Supplementary Figure 1B**), the functions of other eTudor domains lacking an intact aromatic cage remain unclear.

To advance the molecular and mechanistic understanding of *Drosophila* piRNA factors, we aimed to systematically test and characterize direct interactions between them. However, one of the major challenges in studying these proteins using purified proteins and conventional biochemical methods is their structural complexity, as many factors are large multidomain proteins that also contain a substantial degree of predicted disorder. To overcome this challenge, we have previously developed a simple and rapid cell culture-based protein-protein interaction assay, called ReLo, which allows the testing of interactions between structurally complex proteins and specifically detects direct interactions (Salgania et al, 2024). Here, we used ReLo to screen for pairwise interactions between 22 cytoplasmic and mitochondrial surface proteins. As a result, we detected all previously reported interactions for which crystal structures have been determined, as well as several interactions previously identified by co-immunoprecipitation experiments. In addition, we identified previously unknown interactions. To characterize the confirmed and novel interactions at the molecular level, we used a combination of ReLo assays, *in vitro* binding studies, and structural modeling using AlphaFold-Multimer. Here, we present data that shed light on the complexes formed by the TDRD Vreteno and the Yb/TDRD12-related RNA helicases BoYb and SoYb, as well as the RNA helicase Armitage and a heterodimer composed of the mitochondrial proteins Gasz and Daedalus. We also present data describing a multisubunit complex consisting of Tejas (TDRD5), the RNA helicases Vasa (DDX4) and Spindle E (TDRD9), and either Maelstrom or the TDRD Krimper. Together, our ReLo-based screen revealed direct interactions between critical factors in the *Drosophila* piRNA pathway, and we believe that our study will pave the way for future research into the molecular mechanisms underlying piRNA biogenesis, amplification, and function.

## RESULTS

### Pairwise protein-protein interaction (PPI) screening of piRNA pathway proteins

To define direct interactions between proteins essential for piRNA biogenesis and function, we developed the ReLo assay, a rapid and simple cell culture-based PPI method with a colocalization readout (Salgania et al, 2024) (**Figure 1A**). In the ReLo assay, the two proteins of interest are fused to mCherry and mEGFP, respectively, transiently expressed in *Drosophila* S2R+ cells, and their localization is visualized in live cells using confocal fluorescence microscopy. Importantly, one of the constructs is fused to a pleckstrin homology (PH) domain, which directs its localization to the plasma membrane. The colocalization of the second non-membrane anchored protein with the partner at the plasma membrane indicates an interaction. Importantly, we have previously shown that interactions detected by the ReLo assay are highly likely to be direct (Salgania et al, 2024).

**Figure 1.**
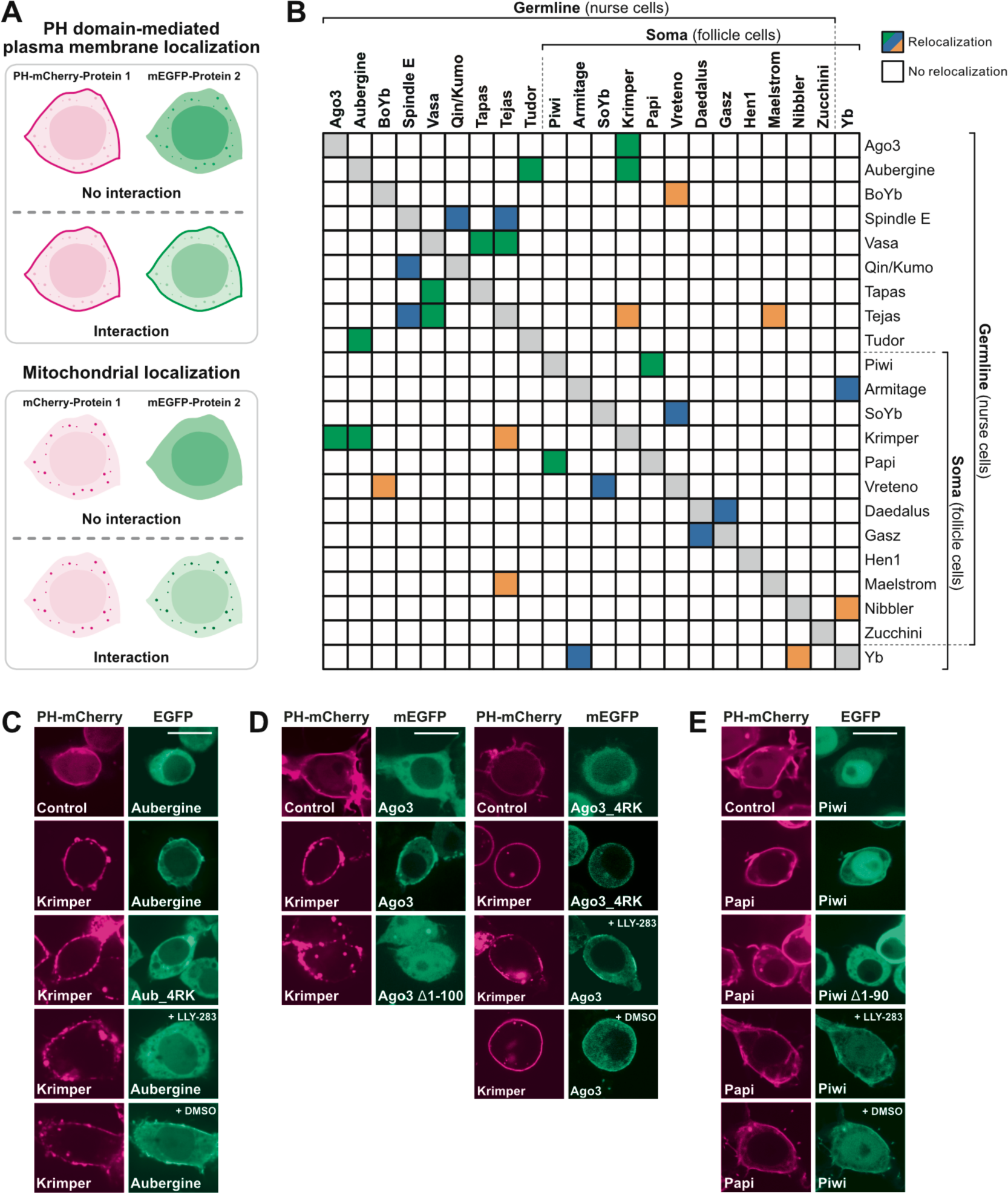
ReLo assay-based PPI screen of cytoplasmic and mitochondrial surface piRNA pathway factors. (A) Principle of the ReLo assay. Two constructs, as indicated, were coexpressed in S2R+ cells, and an interaction between them was inferred if non-membrane anchored protein 2 relocalized to the site of anchored protein 1. Adapted from Figure 1 of Salgania et al. (Salgania et al. 2024) (B) Summary of the PPI screen. Colored rectangles indicate PPIs that were previously characterized based on crystal structures (green), previously observed in co-IPs (blue), or previously unknown (orange). (C) The interaction between Krimper and Aubergine was sDMA-dependent in ReLo assays. The interactions between Krimper and Ago3 (D) and between Papi and Piwi (E) were sDMA-independent. LLY-283 and DMSO concentrations were 0.5 µM and 0.005%, respectively, in (C) and (E) and 1 µM and 0.01%, respectively, in (D). The 4RK mutations in Aub and Ago3 were R11K/R13K/R15K/R17K and R4K/R68K/R70K/R72K, respectively. Scale bars are 10 µm.

Here, we used the ReLo method to perform a pairwise interaction screen of a total of 22 *Drosophila* piRNA factors that are predominantly localized in the cytoplasm or outer mitochondrial membrane *in vivo* (**Figure 1B and Supplementary Figure 2**). To test mitochondrial surface proteins with ReLo, we either used the mitochondrial localization of the proteins without adding the PH domain when testing against non-anchored factors. Alternatively, we used variants of the proteins in which the mitochondrial localization signal (MLS) was deleted when testing against PH-anchored factors (**Figure 1A**). As a result, we detected all previously reported interactions for which crystal structures have been determined, confirmed several interactions previously identified by co-immunoprecipitation experiments, and identified previously unknown interactions (**Figure 1B**). With the exception of the endoribonuclease Zuc and the methyltransferase Hen1, we detected at least one interaction partner for each protein tested (**Figure 1B, Supplementary Figure 2**). Importantly, we only detected interactions that were consistent with previous genetic data. In this study, we focused on the characterization of protein complexes that were previously undescribed or poorly characterized at the molecular level.

### PIWI protein interactions

As a proof of concept, we used ReLo assays to test interactions with PIWI proteins in detail, many of which have been characterized previously. PIWI proteins are known to contain symmetric dimethylated arginine (sDMA) residues within their flexible N-terminal regions (Kirino et al, 2010, 2009), which can be specifically recognized and bound by extended Tudor (eTud) domains present in single or multiple copies in TDRDs (Gan et al, 2019; Handler et al, 2011). The addition of sDMA to PIWI proteins is catalyzed by the protein arginine methyltransferase 5 (PRMT5, a.k.a. Capsuléen in *Drosophila*) (Kirino et al, 2009). By testing and detecting the strictly sDMA-dependent interaction between the PIWI protein Aub and the TDRD Tudor (Liu et al, 2010a; Kirino et al, 2010), we have previously shown that ReLo is suitable for assessing PRMT5-catalyzed sDMA-dependent PPIs (Salgania et al, 2024). Here, we investigated interactions with all three PIWI proteins using the ReLo assay.

In addition to its association with Tudor, Aub has previously been shown to bind to other TDRDs, including Krimper, Tejas, Tapas, and Qin (Patil & Kai, 2010; Patil et al, 2014; Anand & Kai, 2012; Webster et al, 2015; Huang et al, 2021). Of these, we confirmed the binding of Aub to Krimper, but not to the other proteins (**Figure 1C; Supplementary Figure 2C**). Krimper is a critical component that coordinates the Aub/Ago3-driven ping-pong amplification of piRNAs by preventing homotypic Aub-Aub piRNA amplification (Sato et al, 2015; Webster et al, 2015). In ReLo assays, Krimper did not interact with an Aub mutant, in which four methylatable arginine residues in the N-terminus were replaced by lysine residues (R11K/R13K/R15K/R17K, Aub_4RK) and thus do not serve as a substrate for sDMA methylation (Kirino et al, 2009) (**Figure 1C**). Furthermore, when the PRMT5 inhibitor LLY-283 (Bonday et al, 2018) was present in the cell culture medium, an interaction between Krimper and wild-type Aub was not detected, whereas it was detected in the DMSO control experiment (**Figure 1C**). Taken together, these data suggest that the Aub-Krimper interaction is also sDMA-dependent in the ReLo assay, which is consistent with previous reports (Huang et al, 2021; Webster et al, 2015).

The PIWI protein Ago3 was previously reported to bind to the TDRDs Tudor, Krimper, and Tapas (Nishida et al, 2009; Sato et al, 2015; Webster et al, 2015; Huang et al, 2021; Patil et al, 2014). In the ReLo PPI screen, we detected Krimper as the sole interaction partner of Ago3 (**Figure 1D, Supplementary Figure 2**). Deletion of the RG-rich N-terminus of Ago3 (Ago3ϕλ1-100) abolished the Krimper interaction (**Figure 1D**), but it was observed when the Ago3 arginine residues known to be methylated (Nishida et al, 2009) were replaced by lysine residues (R4K/R68K/R70K/R72K; Ago3_4RK) (**Figure 1D**). Furthermore, the PRMT5 inhibitor LLY-283 was unable to inhibit the interaction between Krimper and wild-type Ago3 (**Figure 1D**). Taken together, these data are consistent with previous findings showing that Krimper binds to the N-terminus of Ago3 independently of sDMA modifications (Sato et al, 2015; Webster et al, 2015; Huang et al, 2021).

The *Drosophila* PIWI protein Piwi has been shown to interact with the RNA helicase Armi, and the TDRDs Papi and Qin (Liu et al, 2011; Anand & Kai, 2012; Zhang et al, 2018; Pandey et al, 2017). In our screen, we confirmed Papi but did not detect any other interaction partner of Piwi (**Figures 1B and 1E, Supplementary Figure 2**). Papi has been proposed to associate with all three PIWI proteins (Liu et al, 2011), but we only observed the interaction with Piwi, which is consistent with other previous data (Zhang et al, 2018). There, the Papi-Piwi interaction was shown to depend on both sDMA (R10) and unmethylated arginines (R7, R9, R11) of Piwi and to be mediated by the eTudor domain of Papi (Zhang et al, 2018). Consistent with these data, deletion of the N-terminus of Piwi (Piwi Δ1-90) abolished the Papi interaction (**Figure 1E**). We also tested the effect of the PRMT5 inhibitor LLY-283 and found that it was unable to inhibit the Papi-Piwi interaction (**Figure 1E**), suggesting that this interaction is predominantly sDMA-independent in the ReLo assay.

Taken together, our PIWI interaction tests confirmed all interactions previously characterized using structural information (Liu et al, 2010a; Huang et al, 2021; Zhang et al, 2018), suggesting that with ReLo it should be possible to detect interactions between less well-characterized piRNA factors and interactions that may depend on sDMA modification of one of the partners.

### The Vret - BoYb/SoYb interaction

Next, we investigated in more detail complexes formed by the TDRD Vreteno and the Yb-related proteins BoYb or SoYb (**Figure 2A**), interactions detected in our screen that have been poorly or not characterized at all. *Drosophila* Vreteno is most closely related to mouse TDRD1 (Handler et al, 2011). Vreteno localizes to the nuage in germline nurse cells and to Yb bodies in somatic follicle cells and plays a critical role in piRNA processing in both cell types (Handler et al, 2011; Zamparini et al, 2011); however, its mechanism of action in *Drosophila* is unknown. Vreteno has been suggested to interact with Aub, Ago3, Piwi, Armi, Yb, and the Yb-related protein SoYb (Hirakata et al, 2019; Zamparini et al, 2011; Handler et al, 2011). In our screen, we confirmed SoYb, and also identified BoYb as Vreteno binding partner (**Figures 1B and 2B, Supplementary Figure 2**), two proteins that play a critical role in piRNA biogenesis (Handler et al, 2011). While SoYb is predominantly active in somatic follicle cells, BoYb plays a dominant role in the germline (Handler et al, 2011). It was previously suggested that SoYb binds to Armitage (Hirakata et al, 2019). However, in our pairwise screen did not observe any SoYb or BoYb interaction partners other than Vreteno (**Figure 1B, Supplementary Figure 2**).

**Figure 2.**
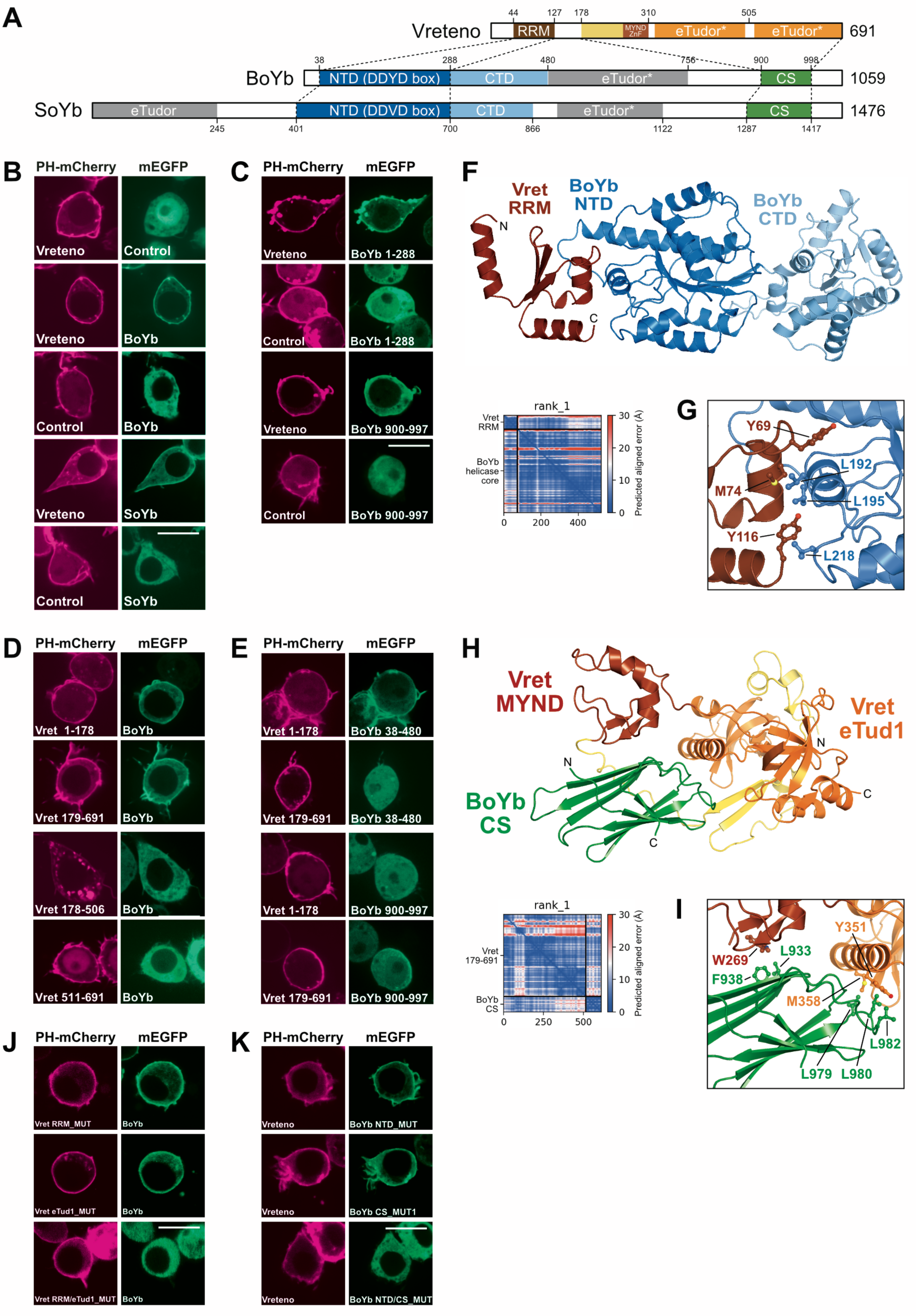
Interaction between Vreteno and BoYb. (A) Domain organization of Vreteno, BoYb and SoYb. (B) Vreteno binds to BoYb and SoYb in ReLo assays. (C) The NTD and the CS domains of BoYb are required for Vret binding. (D) The N- and C-terminal portions of Vret are required for BoYb binding. (E) The N-terminal part of Vret binds the NTD of BoYb and the C-terminal part of Vret binds the CS domain of BoYb. (F) AlphaFold-Multimer-generated structural model of the complex consiting of the Vret-RRM (aa 43-117) and the helicase core of BoYb (aa 38-476). The PAE plot indicates the high confidence of the prediction. (G) Detail of the interface formed by the Vret-RRM and the BoYb core. Residues mutated for subsequent studies are highlighted in ball-and-stick representation. (H) AlphaFold-Multimer-generated structural model of the complex consisting of the C-terminal part of Vret (aa 179-691) with the MYND domain and the eTudor domain bound to the CS domain of BoYb (aa 901-998). The PAE plot shows the high confidence of the model. (I) Detail of the interface formed by the C-terminal part of Vret and the BoYb CS domain. Residues mutated for subsequent studies are highlighted in ball-and-stick representation. (J) Combined mutations in the RRM (RRM_MUT: Y69E/M74E/Y116E) and eTudor (eTud1_MUT: Y351E/M358E) surfaces of Vret are required to abolish the BoYb interaction. (K) Mutations in both the NTD (NTD_MUT: L192E/L195E/L218E) and the CS domain (CS_MUT1: L979E/L980E/L982E) of BoYb are required to abolish the Vret interaction. Scale bars are 10 µm.

Vreteno is composed of an N-terminal RNA recognition motif (RRM), a central MYND-type zinc finger domain, and two C-terminal eTudor domains, both of which lack an intact aromatic cage required for sDMA binding (Handler et al, 2011) (**Figure 2A**). BoYb and SoYb are superfamily 2 RNA helicases with a very similar domain organization (**Figure 2A**) and a helicase core similar to that of DEAD-box RNA helicases. In both proteins, the helicase core is C-terminally followed by an eTudor domain and a ‘CHORD and Sgt1’ (CS) domain, the latter of which is also found in co-chaperones (Rios et al, 2024; Johnson, 2021). In contrast to BoYb, the helicase core of SoYb is N-terminally flanked by an additional eTudor domain. Using ReLo assays, we found that the interaction between Vreteno and SoYb or BoYb maps to two equivalent regions. First, the N-terminal part carrying the RRM domain of Vret (Vret 1-178; Vret-N) bound to the N-terminal RecA-like domain (NTD) of the helicase cores of SoYb and BoYb (**Figures 2C-E, Supplementary Figures 3A-D**). Second, the C-terminal part of Vreteno (Vret 178-691; Vret-C) covering the MYND zinc finger and the two eTudor domains (eTud1 and eTud2) bound to the CS domain of SoYb and BoYb (**Figures 2C-E, Supplementary Figures 3A-D**). In bridging experiments in which PH-mCherry-SoYb was coexpressed with mEGFP-BoYb and non-fluorescent Vreteno, we did not detect any SoYb-BoYb interaction, suggesting that the association of Vreteno with BoYb and SoYb is mutually exclusive (**Supplementary Figure 3E**).

We subjected the two Vret-BoYb interaction modules to structural prediction analysis using AlphaFold-Multimer v.2 (Evans et al, 2021; Mirdita et al, 2022; Jumper et al, 2021) and obtained high-confidence models, as indicated by the low predicted aligned errors (**Figures 2F-I**). In the first model, the Vret RRM domain contacts the face of the NTD of BoYb that is opposite to the position of the C-terminal RecA-like domain (CTD) of the helicase core (**Figure 2F**). In the second model, the CS domain of BoYb contacts both the MYND zinc finger and the eTud1 of Vret-C (**Figure 2H**). For the corresponding Vret-SoYb subcomplexes, we obtained similarly well predicted structural models (**Supplementary Figures 3F and 3G**).

To validate the predicted models, we tested BoYb-Vret surface point mutations in the ReLo assay. When Vret-N carried interface mutations in the RRM domain (RRM_MUT, Y69E/M74E/Y116E) it was unable to interact with BoYb (**Supplementary Figure 3H**). Similarly, when BoYb contained interface point mutations at the NTD (NTD_MUT, L192E/L195E/L218E), it did not interact with Vret-N (**Supplementary Figure 3H**). Taken together, these analyses confirmed the predicted interface formed by the RRM domain of Vret and the NTD of BoYb. Next, we tested mutations located at the BoYb-CS - Vret-C interface. Vret-C carrying mutations in the eTud1 domain (eTud1_MUT, Y351E/M358E) as well as BoYb carrying surface mutations in the CS domain facing the eTud1 domain of Vret (CS_MUT1, L979E/L980E/L982E) and those facing the MYND domain (CS_MUT2, L933E/F938E) prevented the interaction (**Supplementary Figure 3I)**. These data support the predicted structural model of the complex formed between the CS domain of BoYb and Vret-C. We also tested the individual interface mutations in the context of the full-length protein and found that they only inhibited the interaction when they were combined: while the combined mutations in Vreteno (RRM_MUT and eTud1_MUT) inhibited the interaction with BoYb, the individual sets of mutations did not (**Figure 2J**). Similarly, the combined mutations in BoYb (NTD_MUT and CS_MUT1) inhibited binding to Vreteno, but the individual sets of mutations were not sufficient (**Figure 2K**).

In summary, our data suggest that the Vreteno - BoYb and Vreteno - SoYb complexes form through two independent binding interfaces. Subjecting the two complexes to a DALI search (Holm & Rosenström, 2010) did not yield any previously known similar complex structure, suggesting that our analyses have revealed two novel interaction modules.

### The Gasz - Daed interaction

Next, we characterized the complex formed by Daedalus (Dead) and Gasz, proteins localized to the outer mitochondrial membrane that, as a complex, have been proposed to be required for the translocation of Armi and Piwi to mitochondria (Olivieri et al, 2012; Munafò et al, 2019; Yan et al, 2002; Zhang et al, 2016). In our pairwise screen, we observed the Gasz-Daed interaction, but did not identify any other protein that bound to either component individually (**Figures 1B and 3A-C, Supplementary Figure 2**).

**Figure 3.**
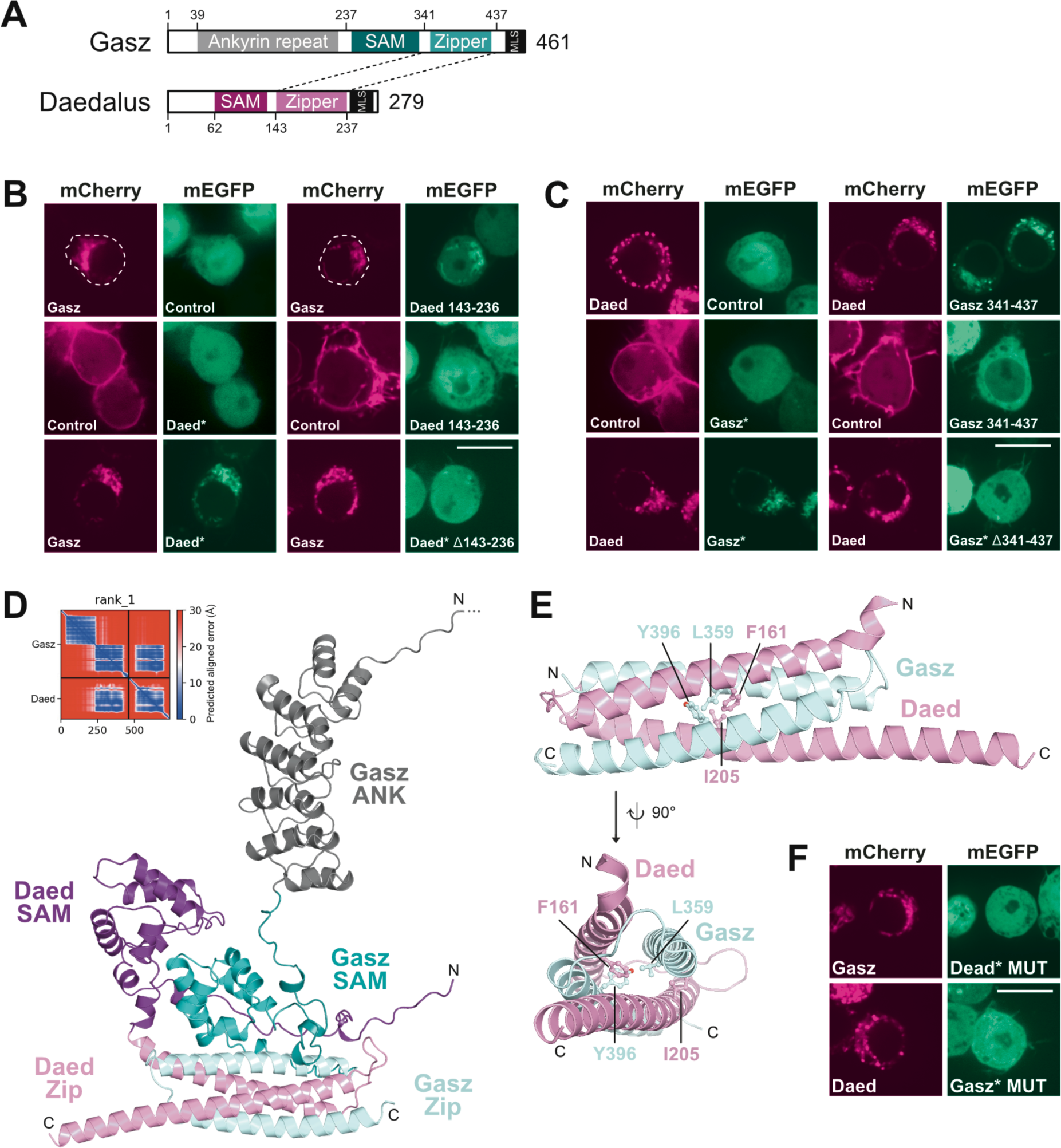
Interaction between Gasz and Daed. (A) Domain organization of Gasz and Daed. (B, C) Daed bound to Gasz in ReLo assays. The interaction required the Zipper domain of Daed (B) and Gasz (C). An asterisk indicates that the protein (additionally) lacks the C-terminal MLS, corresponding to aa 438-461 in Gasz and aa 237-279 in Daed. (D) AlphaFold-Multimer-generated structural model of the Daed-Gasz complex (missing the MLSs). The PAE plot indicates the high confidence of the prediction. (E) Detail of the interface formed by the Zipper domains of Gasz and Daed. Residues mutated for subsequent studies are highlighted in ball-and-stick representation. (F) Mutations in the Zipper interface prevented the formation of the Gasz-Daed complex. Scale bars are 10 µm.

Gasz and Daed share a similar domain organization consisting of a sterile-α-motif (SAM), followed by a leucine zipper (Zip) and a C-terminal transmembrane α-helix that serves as a mitochondrial localization signal (MLS) (Munafò et al, 2019; Yan et al, 2002). Gasz contains an additional uncharacterized ankyrin repeat (ANK) domain at the N-terminus (**Figure 3A**) (Yan et al, 2002). Using ReLo assays, we mapped the Gasz-Daed interaction to their respective Zip domains (**Figures 3B and 3C, Supplementary Figure 4**), which is consistent with previous co-IP data (Munafò et al, 2019). Using AlphaFold-Multimer v2 (Evans et al, 2021; Jumper et al, 2021; Mirdita et al, 2022), we generated a structural model of the Gasz - Daed heterodimer, in which the Zip domains of Gasz and Daed are extensively intertwined (**Figure 3D**). The model also shows extensive contact between the respective SAM domains. Since the SAM domains alone were neither necessary nor sufficient for Gasz-Daed interaction in ReLo assays (**Supplementary Figure 4**), the SAM-SAM contact may be weak and established only after the Zip heterodimer has formed. Mutation of two critical Zip interface residues in either Gasz (Gasz MUT; L359E/Y396E) or Daed (Daed MUT; F161E/I205E), abolished the interaction (**Figures 3E and 3F**), confirming the predicted structural model of the Zip heterodimer.

### The Gasz-Dead-Armi complex

Previously Daed and Gasz were proposed to recruit the RNA helicase Armi (**Figure 4A**) to mitochondria (Munafò et al, 2019). Since we did not detect any proteins other than Daed and Gasz that bound to Gasz or Daed in our pairwise screen, we asked whether Gasz and Daed bind to Armi as a complex in an experimental setup in which mCherry-Gasz was coexpressed with both mEGFP-Armi and non-fluorescent Daed. Indeed, an interaction between Gasz and Armi was detected in the presence of Daed (**Figure 4B**). When the ternary interaction was tested using constructs containing point mutations located at the Gasz-Daed interface (Daed-MUT or Gasz-MUT), no interaction was observed (**Figure 4B**), further suggesting that Armi binding requires a complex formed by Gasz and Dead.

**Figure 4.**
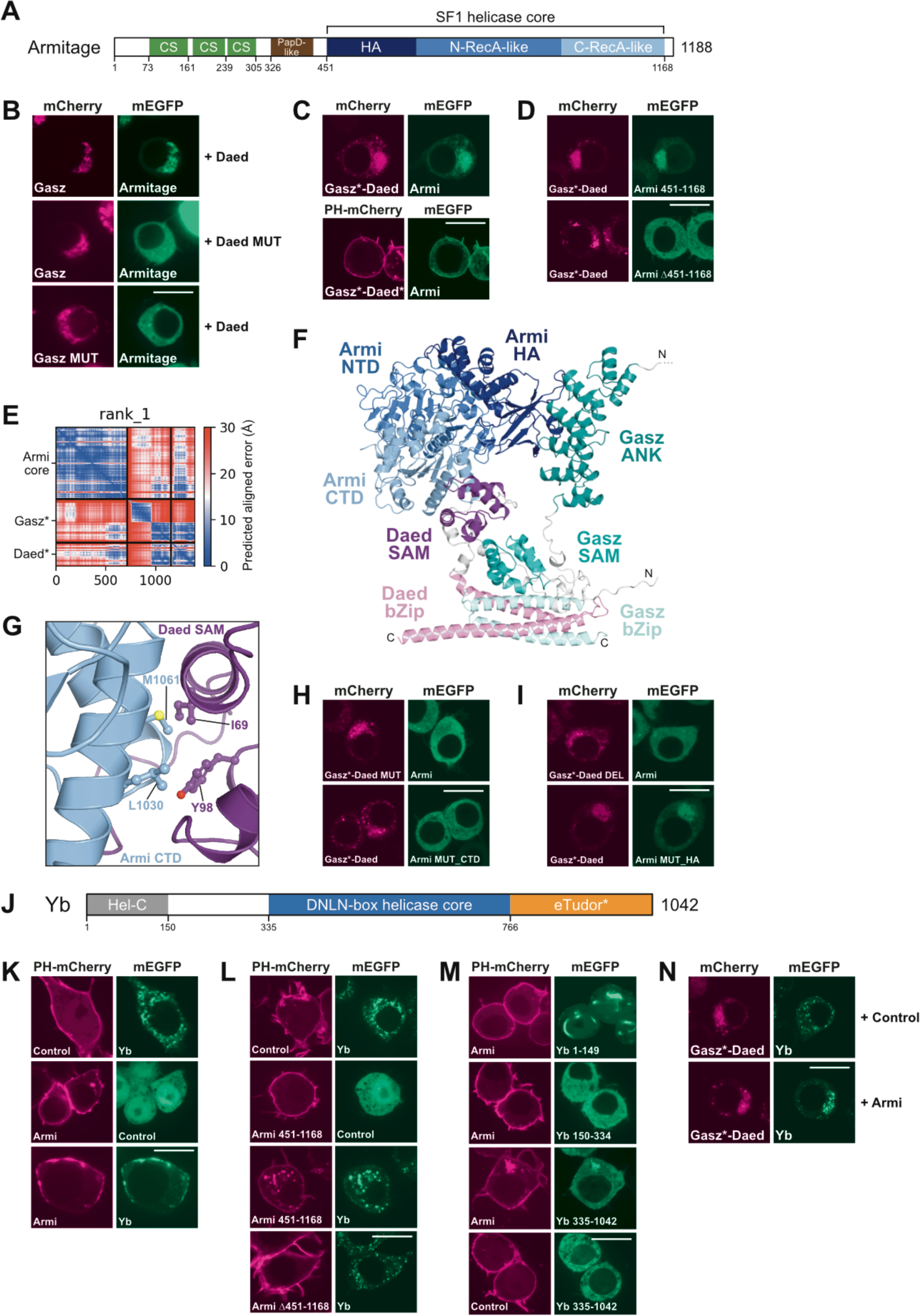
Interaction between Armi and the Gasz-Daed complex. (A) Domain organization of Armi. (B) Armi binds to Gasz in the presence of Daed, but not in the presence of Daed carrying the mutation (Daed MUT: F161/I205) that abolishes Gasz-Daed complex formation. Armi does not bind to Gasz MUT (L359/Y396E) in the presence of Daed. (C) A Gasz*-Daed fusion lacking the Gasz-MLS recruits Armi to mitochondria. A Gasz-Daed fusion carrying the PH domain at the N-terminus and lacking both MLS (Gasz*-Daed*) recruited Armi to the plasma membrane. (D) The Gasz*-Daed fusion binds to the helicase core of Armi (aa 451-1168). (E) PAE plot showing the high confidence of the prediction in (F). (F) AlphaFold-Multimer-generated structural model of the Daed-Gasz complex (lacking the MLSs) bound to the helicase core (aa 451-1168) of Armi. (G) Detail of the interface formed by the CTD of Armi and the SAM domain of Daed. Residues mutated for subsequent studies are highlighted in ball-and-stick representation. (H) Mutations in the CTD-SAM interface prevented the formation of the Gasz-Daed-Armi complex. The Gasz-Daed MUT is I69E/Y98E located in the Daed part and the Armi MUT_CTD is L1030E/M1061E. (I) Deletion of the Gasz Ankyrin repeat (aa 1-236) from the Gasz-Daed fusion (DEL) abolished Armi binding. Mutations at the HA surface of Armi (MUT_HA, W592E/D631A) did not prevent binding to the Gasz-Daed fusion. (J) Domain organization of Yb. (K) Yb interacts with Armi in ReLo assays. (L) The Armi helicase core (aa 451-1168) is necessary and sufficient for Yb binding. (M) Armi binds to the C-terminal portion (aa 335-1042) of Yb, which includes the helicase core and the eTudor domain. (N) Yb binds to the Gasz-Daed fusion in the presence of Armi. The asterisk (*) indicates when the mitochondrial localization signal was deleted from the protein. Scale bars are 10 µm.

Armi is an ATP-dependent enzyme that belongs to the superfamily 1 of RNA helicases, and is predicted to be structurally similar to Upf1 (Pandey et al, 2017; Ishizu et al, 2019; Yamashiro et al, 2020). The helicase core is N-terminally flanked by an array of three cold shock (CS) domains, followed by a PapD-like fold (**Figure 4A**). To map the region of Armi that binds to the Gasz - Daed complex, we used a two-component setup in which the various mEGFP-Armi constructs were cotransfected with a single construct consisting of mCherry-tagged Gasz lacking the MLS and fused C-terminally to Daed (mCherry-Gasz-Daed). This latter construct localized to mitochondria through the MLS of Daed and showed interaction with full-length Armi in ReLo assays, suggesting that it is functional (**Figure 4C**). We found that Armi also interacted with the Gasz-Dead fusion when tested for relocalization to the plasma membrane, suggesting that the interaction *per se* is independent of mitochondrial localization (**Figure 4C**).

In mapping experiments, the Armitage helicase core (Armi 451-1168) was necessary and sufficient for binding to the Gasz-Daed fusion (**Figure 4D, Supplementary Figures 5A and 5B**). Using AlphaFold-Multimer prediction of a complex composed of Gasz and Daed, both lacking the MLS, and the Armi helicase core, we obtained a high confidence structural model showing that the SAM domain of Daed contacts the C-terminal RecA-like domain of the Armi core (**Figures 4E and 4F**). In addition, a less confident prediction is observed for the Ankyrin repeat of Gasz contacting the helicase-associated (HA) domain of the Armi core. The Armi - Daed-SAM interface was validated by interfering point mutations on both sides, which abolished the interaction (**Figures 4G and 4H**). While mutations in the Gasz - Armi interface did not interfere with complex formation (**Figure 4I, Supplementary Figure 5C**), deletion of the Ankyrin repeat from the Gasz-Daed fusion abolished the interaction with Armi (**Figure 4I**), suggesting that the Ankyrin repeat contributes to ternary complex formation but the interface remains unclear.

In addition to the Gasz-Daed complex, Armitage has been shown to associate with the PIWI proteins Ago3 and Piwi and the TDRDs Vreteno and Yb in co-IP experiments (Huang et al, 2014; Handler et al, 2011; Hirakata et al, 2019; Zamparini et al, 2011; Haase et al, 2010; Olivieri et al, 2010; Pandey et al, 2017; Saito et al, 2010a). Of these, we only detected the Yb-Armi interaction in pairwise ReLo tests (**Figures 1B and 4K, Supplementary Figure 2**). Yb is composed of an N-terminal Hel-C domain, which shows structural similarity to C-terminal RecA-like helicase domains, a central DEAD-box-like RNA helicase core, and a C-terminal eTudor domain (**Figure 4J**). The Hel-C domain (aa 1-150) has been discussed to promote self-association (Hirakata et al, 2019). Testing for potential dimer, trimer, or tetramer formation of the Hel-C domain using AlphaFold-Multimer did not result in a high confidence model prediction (**Supplementary Figure 5D**), and the molecular basis for the proposed self-association of this domain remains unclear. Using ReLo assays, we mapped the Armi-Yb interaction to the helicase core of Armi and the C-terminal half of Yb, which covers the helicase core and the eTudor domain (**Figures 4L and 4M; Supplementary Figure 5E**). We did not obtain a high-confidence model of the Armi-Yb complex from an AlphaFold-Multimer prediction run (Jumper et al, 2021; Evans et al, 2021; Mirdita et al, 2022), so we did not further characterize this interaction. Nevertheless, we tested whether Yb can associate with the Armi-Gasz-Daed complex and found that it does in ReLo assays (**Figure 4N**), suggesting that Yb and the Gasz-Daed complex contact different surfaces of the Armi helicase core for binding.

### The Tejas - Spindle E interaction

Next, we examined protein interactions with Tejas (the *Drosophila* ortholog of TDRD5), a central component of nuage that was previously shown to be required for ping-pong amplification of piRNAs (Patil & Kai, 2010; Patil et al, 2014). Tejas contains an N-terminal eLOTUS domain and a C-terminal eTudor domain of unknown function (**Figure 5A**). Tejas interacts with the ATP-dependent DEAD-box RNA helicase Vasa through its eLOTUS domain, and eLOTUS domains have been shown to stimulate the ATPase activity of Vasa (Jeske et al, 2017). The precise function of Tejas in this pathway remains unclear. Tejas has also been suggested to associate with the TDRDs Tapas (*Drosophila* ortholog of TDRD7) and Spindle E (a.k.a. Homeless; *Drosophila* ortholog of TDRD9) (Patil & Kai, 2010; Patil et al, 2014; Lin et al, 2023), and we confirmed Tapas (**Figure 1B, Supplementary Figure 2P**) and Spindle E as Tejas binding partners (**Figures 1B and 5B, Supplementary Figure 2**). Because we did not obtain a high confidence structural model for the Tejas - Tapas complex, we did not further characterize this interaction.

**Figure 5.**
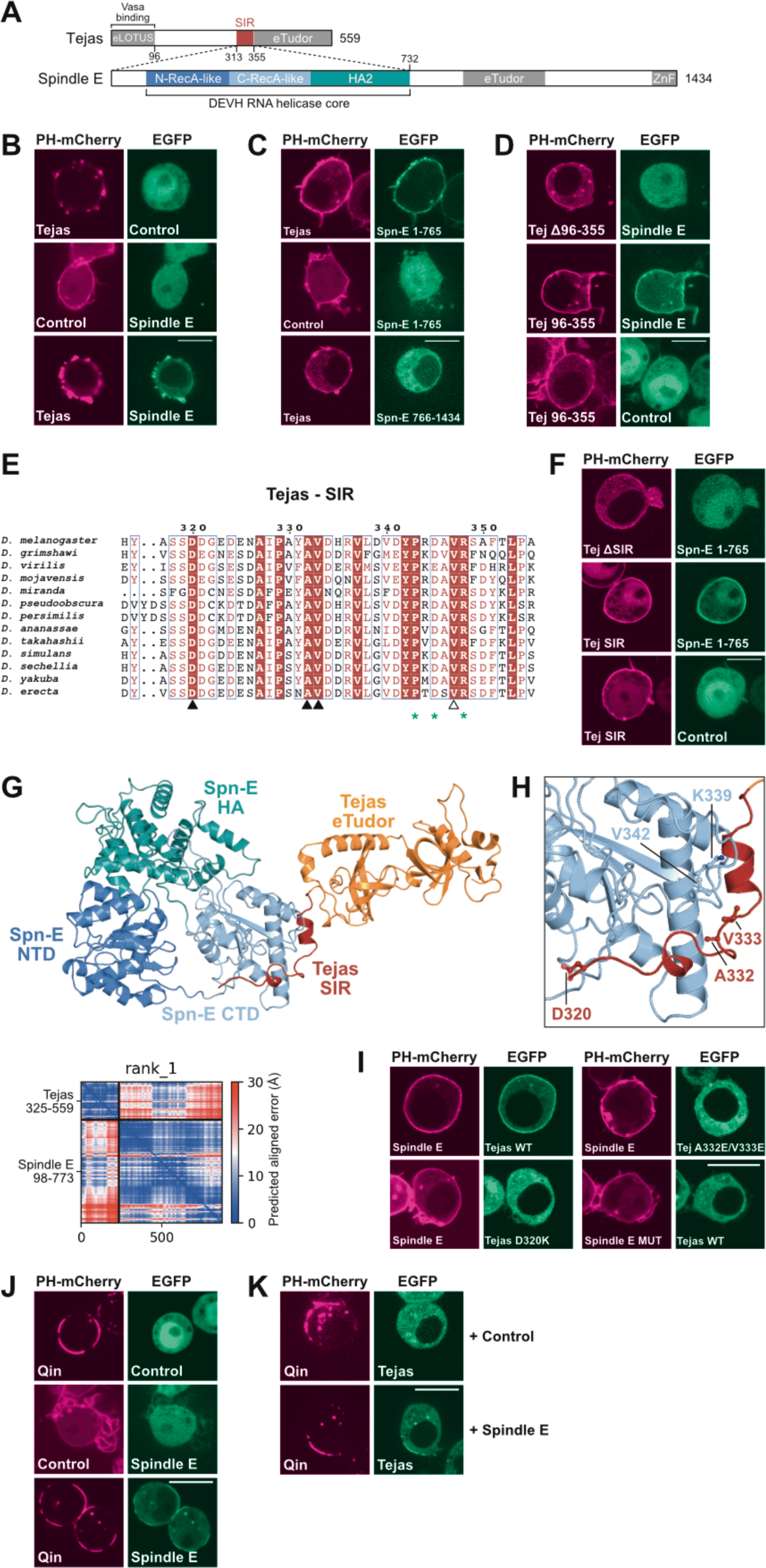
The Interaction between Tejas and Spindle E. (A) Domain organization of Tejas and Spindle E. (B) Tejas interacts with Spindle E in ReLo assays. (C, D) The interaction is mediated by the N-terminal portion of Spindle E (C) and the predominantly unstructured region of Tejas (D). (E) Multiple sequence alignment showing the SIR, which is the most conserved region of the predominantly unstructured region of Tejas and extends N-terminally of the eTudor domain. (F) The SIR of Tejas is necessary and sufficient for binding to Spindle E. (G) AlphaFold-Multimer-generated structural model of the complex formed by the helicase core of Spindle E (aa 98-773) and the C-terminal part of Tejas comprising the SIR and the eTudor domain (aa 325-559). The PAE plot indicates the high confidence of the prediction. (H) Detail of the interface formed by the Tejas-Spindle E complex. Residues mutated for subsequent studies are highlighted in ball-and-stick representation. (I) Mutations in the predicted interface, as indicated, prevented formation of the Tejas-Spindle E complex. Spindle E MUT is K339/V342E. Scale bars are 10 µm. (J) Spindle E bound to Qin in ReLo assays. (K) Qin bound to Tejas in the presence of Spindle E. Scale bars are 10 µm.

Spindle E is critical for the ping-pong amplification of piRNAs (Malone et al, 2009; Gillespie & Berg, 1995), but its molecular function in this process in *Drosophila* remained unclear. Spindle E belongs to the DEVH-box family of RNA helicases with a helicase core flanked C-terminally by an eTudor domain and a zinc finger (ZnF) (**Figure 5A**). Using ReLo assays, the Tejas-Spindle E interaction was mapped to the N-terminal half of Spindle E containing the helicase core and to a short, conserved region of Tejas that extends N-terminally from the eTudor domain, which we termed the Spindle E-interacting region (SIR) (aa 315-355) (**Figures 5C-F, Supplementary Figure 6A**). Tejas binding to Spindle E is independent of mutations in the putative ATP binding pocket of Spindle E (**Supplementary Figure 6B**). Using AlphaFold-Multimer-v3 (Mirdita et al, 2022; Evans et al, 2021; Jumper et al, 2021) a high confidence structural model of the Tejas-Spindle E complex was predicted, in which the SIR contacts the C-terminal RecA-like domain of the helicase core of Spindle E (**Figure 5G and 5H**). We validated the Tejas-Spindle E interface by mutagenesis of critical interface residues of Tejas (D320K; A332E/V333E) and Spindle E (Spindle E MUT; K339E/V342E) using ReLo assays (**Figure 5I)**. The Tejas mutation V347E had no effect on the Tejas - Spindle E interaction in ReLo assays (**Figure 5I**), suggesting that the SIR can be refined to aa 315-335 of Tejas. An identical structural model of the Tejas-Spindle E complex was also recently reported using AlphaFold2 (Lin et al, 2023). In addition to Tejas, Spindle E has been proposed to interact with the TDRDs Tapas (Patil et al, 2014) and Qin (TDRD4) (Anand & Kai, 2012) and with the RNA helicase Vasa (Pek & Kai, 2011), of which we could only confirm Qin as an additional Spindle E interaction partner (**Figures 1B and 5J, Supplementary Figure 2**). As we did not obtain a high confidence structural model from an AlphaFold-Multimer run, we did not investigate this interaction in more detail. Nevertheless, we tested whether Spindle E could bind to Tejas and Qin simultaneously and found that it did in ReLo bridging assays (**Figure 5K**).

### Tejas binds to Maelstrom or Krimper through its eTudor domain

Our ReLo screen also identified two previously unknown Tejas binding partners, Maelstrom and Krimper. Maelstrom functions in the nucleus in piRNA-dependent and -independent repression of transcription (Chang et al, 2019; Sienski et al, 2012; Klenov et al, 2011; Onishi et al, 2020), but is also found in the perinuclear nuage (Findley et al, 2003; Lim & Kai, 2007; Sienski et al, 2012). A cytoplasmic molecular function for *Drosophila* Maelstrom has not been described. Maelstrom is composed of an N-terminal high mobility group (HMG) box, which in the mouse Maelstrom ortholog binds to RNA *in vitro* (Genzor & Bortvin, 2015), a central Mael domain with a nuclease fold (Matsumoto et al, 2015; Chen et al, 2015), and a C-terminal mostly unstructured region of unknown function (**Figure 6A**). Using ReLo and GST pull-down assays, we mapped the Tejas-Maelstrom interaction to the eTudor domain of Tejas and to the C-terminal region of Maelstrom (**Figures 6B, Supplementary Figures 7A and 7B**). Using AlphaFold-Multimer-v2 (Evans et al. 2021; Jumper et al. 2021; Mirdita et al. 2022), a high-confidence structural model of the complex was predicted, revealing that the most C-terminal part of Maelstrom (aa 413-459), which we termed the Tejas-interacting region (TIR), forms an α-helix that binds to the eTudor domain of Tejas (**Figure 6C**). Interestingly, in this complex, the Mael-TIR does not occupy the canonical peptide-binding cleft formed by both the Tudor domain and the SN fold of the eTudor domain (**Supplementary Figure 1B**), but instead contacts an exposed surface of the SN fold. Deletion of the TIR from Maelstrom abolished the Tejas interaction in GST pull-down and ReLo assays (**Figures 6D and 6E**). The Tejas-Maelstrom interface was further validated by testing mutant variants of Tejas (Tejas MUT; V530E/V532E) and Maelstrom (Mael MUT; V443E/F448E/V450E) in ReLo and GST pull-down assays, which prevented the Tejas-Maelstrom interaction (**Figure 6E and 6F**).

**Figure 6.**
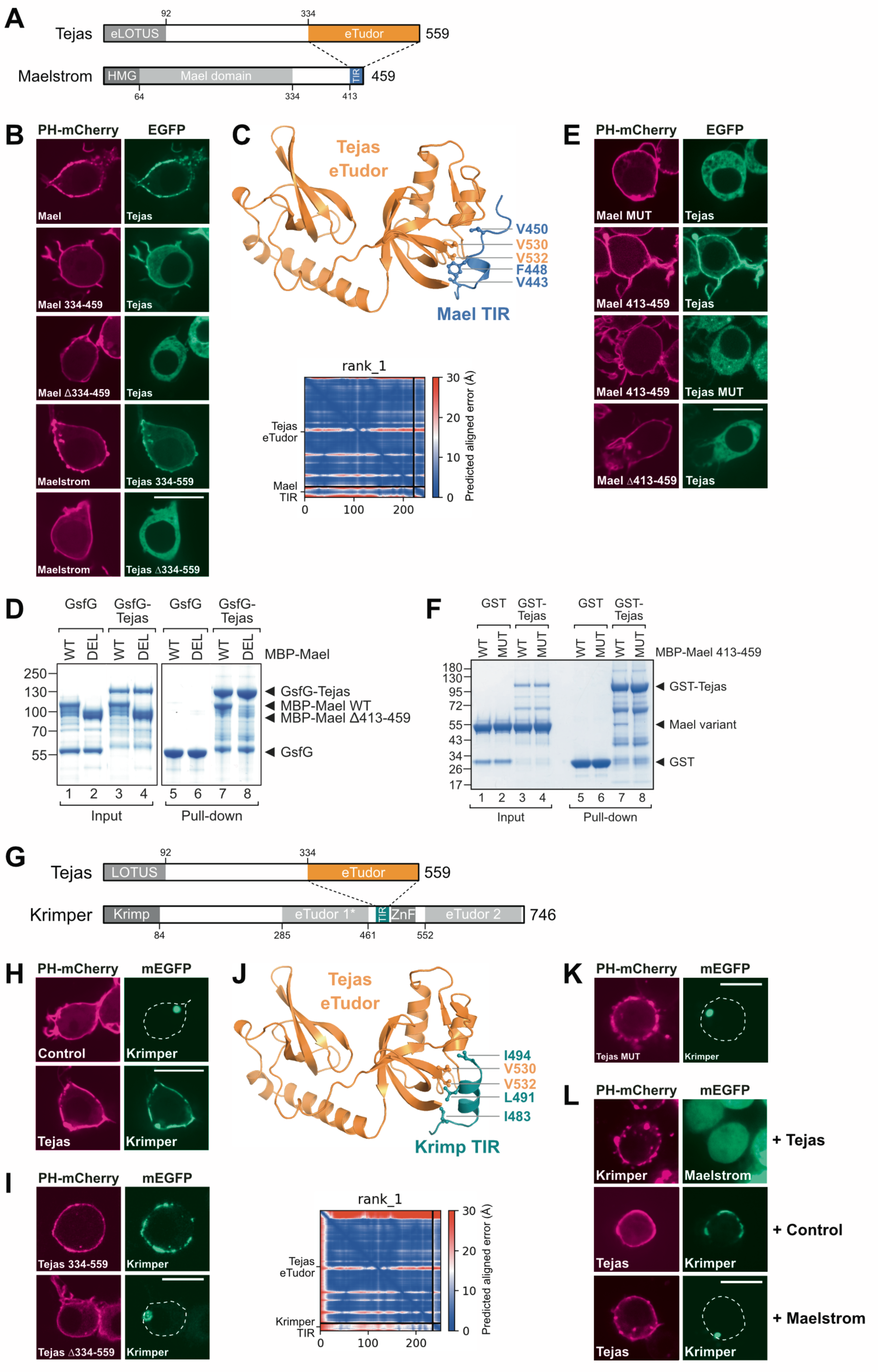
Interaction between Tejas and Maelstrom or Krimper. (A) Domain organization of Tejas and Maelstrom. (B) Tejas interacts with Maelstrom in ReLo assays and the interaction is mediated by the C-terminal region of Maelstrom (aa 334-459) and the eTudor domain of Tejas (aa 334-559). (C) AlphaFold-Multimer-generated structural model of the complex formed by the eTudor domain of Tejas and the Tejas-interacting region (TIR) of Maelstrom. Residues mutated for subsequent studies are highlighted in ball-and-stick representation. The PAE plot indicates the high confidence of the prediction. (D) GST pull-down assay showing binding of GST-superfolder-GFP (GsfG)-Tejas to full-length MBP-Maelstrom but not to Maelstrom lacking aa 413-459. (E) Mutations in the predicted interface prevent Tejas-Maelstrom complex formation. (F) GST pull-down assay showing binding of GST-Tejas to MBP-fused Mael C-terminus (aa 413-459) but not to a mutated version of it. (G) Domain organization of Tejas and Krimper. (H) Tejas interacts with Krimper in ReLo assays. (I) The Krimper interaction is mediated by the eTudor domain of Tejas (aa 334-559). (J) AlphaFold-Multimer-generated structural model of the complex formed by the eTudor domain of Tejas and the Tejas-interacting region (TIR) of Krimper. Residues mutated for subsequent studies are highlighted in ball-and-stick representation. The PAE plot indicates the high confidence of the prediction. (K) Mutations in the predicted Tejas interface prevented Tejas-Krimper complex formation. (L) In ReLo assays, Tejas did not bridge Krimper and Maelstrom (top panels). The Tejas-Krimper interaction is abolished by coexpression of Maelstrom (middle and lower panels). Scale bars are 10 µm.

Krimper is the second previously unknown Tejas binding partner, consisting of an N-terminal coiled-coil (Krimp) domain and two C-terminal eTudor domains (eTud1 and eTud2) (**Figure 6G**), of which eTud2 binds to Aub in an sDMA-dependent manner and eTud1 lacks an intact aromatic cage and binds to Ago3 in an sDMA-independent manner (Huang et al. 2021; Sato et al. 2015; Webster et al. 2015) (see also **Figures 1B-D**). Between the two eTudor domains, Krimper harbors a zinc finger (ZnF) domain of unknown function. The N-terminal Krimp domain has been proposed to mediate homodimerization (Webster et al. 2015) and indeed, a high-confidence structural model of the Krimp domain dimer was predicted using AlphaFold-Multimer (**Supplementary Figure 7C**).

Domain mapping experiments using ReLo assays revealed that Krimper binds to the eTudor domain of Tejas (**Figure 6I and Supplementary Figure 7D**), the same domain bound by Maelstrom. We attempted to map the region in Krimper that binds to Tejas. However, the results were inconclusive. On the one hand, none of the individual protein domains of Krimper bound to Tejas (**Supplementary Figure 7E**), suggesting that more than one weak binding site in Krimper contributes to the Tejas interaction. On the other hand, we obtained ambiguous results from the Krimper domain deletion experiments, likely due to the different subcellular localization of the Krimper deletion constructs. In S2R+ cells, Krimper localized as a large granule, which was previously observed in OSCs and named Krimp body (Olivieri et al. 2012; Sato et al. 2015). Some Krimper deletion constructs also showed granular localization (KrimpΔ84-284, KrimpΔ285-461, and KrimpΔ462-552), but Krimper with a deletion in the Krimp domain (KrimpΔ1-83) or in the eTud2 domain (KrimpΔ551-746) localized ubiquitously (**Supplementary Figure 7F**). When we performed Tejas interaction assays with the individual Krimper deletions, we observed that only the deletions that form a single granule by themselves (KrimpΔ84-284, KrimpΔ285-461) relocalized with Tejas, while the deletions that do not form a single granule did not interact (**Supplementary Figure 7G**). These data suggest that either the Krimp domain and the stretch covering the ZnF and eTud2 domains are all required for Tejas binding, or the Tejas-Krimper interaction is weak and is only observed when Krimper is locally concentrated by accumulating in a single granule.

Nevertheless, we subjected the Tejas and Krimper protein sequences to AlphaFold-Multimer prediction runs and obtained a high confidence model (**Figure 6J**) in which the eTudor domain of Tejas interacts with a predicted α-helix (spanning aa 483-494) of Krimper located between its two eTudor domains (**Figure 6G**). The binding mode observed in the Tejas-Krimper complex is equivalent to that of the Tejas-Maelstrom complex (compare with **Figure 6C**). ReLo assays revealed that the Tejas mutations that disrupted the Maelstrom interaction (Tejas MUT; V530E/V532E) also disrupted the Krimper interaction (**Figure 6K**). However, deletion or mutation of the putative Tejas interaction region (TIR) of Krimper (KrimperΔ483-494 or Krimper MUT: I483E/L491E/I494E) did not disrupt the interaction with Tejas (**Supplementary Figure 7H**), suggesting either that the predicted TIR is incorrect or that additional regions in Krimper contribute to Tejas binding. Thus, the exact region(s) in Krimper that contact Tejas remain unclear from the experiments. Additional ReLo bridging experiments showed that Krimper and Maelstrom do not simultaneously bind to Tejas, but instead compete for Tejas binding in a competition experiment (**Figure 6L**). These observations are consistent with the finding that the same surface on the eTudor domain of Tejas is occupied by Maelstrom or Krimper.

### Tejas and Krimper mediate an interaction of Vasa with Aubergine

So far, we have described and mapped the pairwise interactions between Tejas and Spindle E, Maelstrom, and Krimper. We have also previously characterized the interaction between Vasa and the eLOTUS domain of Tejas (Jeske et al. 2017). Additional studies of the eLOTUS domain of the Oskar protein revealed that it binds to the open but not the closed conformation of Vasa (Jeske et al. 2017; Salgania et al. 2024), and we hypothesized a similar conformation-dependent binding preference for the eLOTUS domain of Tejas. The core of DEAD-box helicases consists of two RecA-like domains that have different orientations relative to each other, depending on whether ATP and RNA are bound or not. In a substrate-unbound form, the helicase core adopts an open conformation that closes upon substrate binding. The open or closed conformational state of a DEAD-box helicase core can be stabilized by specific mutations: The K295N point mutation of Vasa (Vasa-open) abolishes ATP binding and consequently RNA binding, thus maintaining the helicase core in an open conformation; the E400Q point mutation of Vasa (Vasa-closed) allows ATP and RNA binding and subsequent ATP hydrolysis, but prevents dissociation of the ATP hydrolysis products and RNA, thus locking the helicase core in a closed form (Xiol et al. 2014; Walker et al. 1982; Gorbalenya et al. 1988). In ReLo assays, we indeed observed Tejas interacting with Vasa-open but not with Vasa-closed (**Figure 7A**), suggesting a similar binding preference as observed for Oskar. Tejas did not relocalize with Vasa-WT, suggesting that Vasa-WT is in a closed conformation in cells at steady state. We therefore used Vasa-open in all subsequent interaction assays with Tejas.

**Figure 7.**
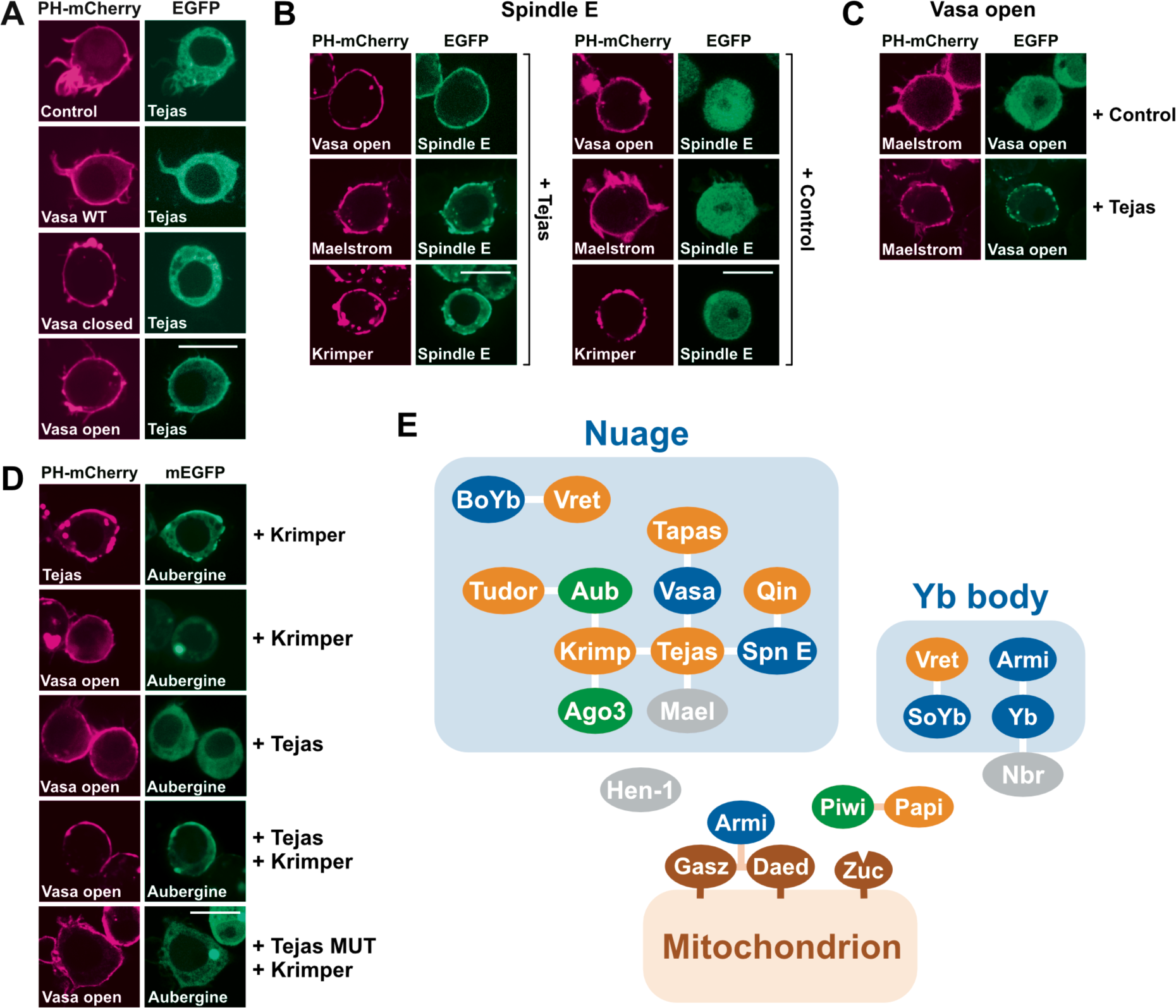
The Tejas interaction network. (A) Tejas relocalized to Vasa only when Vasa carried an ATP-binding mutation that kept it in an open conformation (Vasa-open). (B) Spindle interacted with Vasa-open, Maelstrom or Krimper when Tejas was coexpressed. (C) Maelstrom interacted with Vasa-open when Tejas was coexpressed. (D) Tejas interacted with Aubergine when Krimper was coexpressed (top panels). Vasa-open interacted with Aubergine when both Tejas and Krimper were coexpressed. Coexpression of Krimper and a Tejas mutant that abolishes Krimper binding (Tejas MUT, V530E/V532E) did not mediate the Vasa-open - Aub interaction. (E) Summary of interactions observed in the ReLo PPI screen. Scale bars are 10 µm.

Next, we asked whether some of the Tejas interaction partners could simultaneously bind to Tejas to form a multi-subunit complex, and we performed bridging experiments in which three constructs were coexpressed. In the presence of Tejas, Spindle E relocalized with Vasa-open, Maelstrom, and Krimper (**Figure 7B**), suggesting that the Tejas-Spindle E complex can simultaneously associate with any other individual Tejas partner. In addition, we observed that Vasa-open colocalized with Maelstrom in the presence of Tejas (**Figure 7C**), suggesting that Maelstrom binding does not interfere with the Tejas - Vasa interaction, but that a ternary complex can form. We did not test Krimper in this assay because this protein is only weakly associated with Vasa, and ReLo is not a quantitative assay from which we could conclude that the Krimper - Vasa interaction is more stable in the presence of Tejas. Nevertheless, the data taken together suggest that Tejas can physically assemble a multi-subunit complex containing Spindle E, Vasa, and Maelstrom (or possibly alternatively Krimper).

In our interaction screen, Krimper bound not only to Tejas, but also to Vasa, Aub and Ago3 (**Figure 1**). In bridging experiments, we observed that Krimper was able to mediate the binding of Tejas to Aub (**Figure 7D**), suggesting that Tejas and Aub have distinct binding sites on Krimper and thus Tejas is unlikely to bind to the eTudor2 domain of Krimper. Krimper was unable to mediate an interaction between Vasa and Aub (**Figure 7D**). However, when both Krimper and Tejas were coexpressed, an interaction of Vasa with Aub or Ago3 was observed. Taken together, these data provide insight into a PPI network that can physically assemble around Aub, a central player in the ping-pong amplification step.

## DISCUSSION

Using ReLo, a PPI assay recently established in our lab (Salgania et al. 2024), we present data from a systematic pairwise PPI screen of 22 *Drosophila* cytoplasmic and mitochondrial surface piRNA factors (**Figure 7E**). We have previously shown that PPIs detected by ReLo are most likely direct and not mediated by proteins (or RNA) present in the cells (Salgania et al. 2024). Therefore, while the ReLo assay may not provide information about PPIs that depend on the presence of RNA, such as piRNAs or target mRNAs, the detection of direct interactions is advantageous for the study of piRNA pathway-associated protein complexes by biochemical and structural biology approaches. Indeed, our study combined with AlphaFold structural modeling provides new molecular insights through the in-depth characterization of six previously uncharacterized complexes. It is likely that our methodological approach can be easily adapted to study protein interaction networks of other biological processes in any organism.

### The Gasz-Daed-Armi complex

While several previous lines of evidence suggested complex formation between Gasz and Dead (Munafò et al. 2019), our study revealed how the Zip domains (and potentially also the SAM domains) of Gasz and Dead are involved in heterodimer formation, and we identified specific point mutations that abolish the interaction. Such mutations may be useful for future *in vivo* studies using a transgenic approach.

Structural and experimental data suggest that the SAM domain of Daed contacts the N-terminal RecA-like domain of Armi. Out of curiosity, we performed an AlphaFold 3 prediction run (Abramson et al. 2024) of the Armi-Daed complex bound to a dsRNA oligo containing a single-stranded 5’ overhang, previously used in dsRNA unwinding assays (Ishizu et al. 2019). We obtained a high-confidence model that allows one to imagine where the dsRNA portion might enter and where the ssRNA portion might exit the RNA binding cleft of Armi (**Supplementary Figure 5F**). Interestingly, near the exiting single-strand, we observed a high amount of positively charged amino acid residues emanating from a region of Dead that is N-terminal to the SAM domain (**Supplementary Figure 5F**). Although this basic stretch is not predicted to be highly structured, it is conserved, also considering the similarity of the chemical properties of the residues (**Supplementary Figure 5G**), suggesting that this stretch might be a functional region of Daed possibly involved in RNA binding. It is therefore tempting to speculate that the Gasz-Dead complex might serve to selectively recruit those Armi molecules to mitochondria that are loaded with RNA, a hypothesis that remains to be tested.

### Yb body assembly

Yb bodies are follicle cell-specific perinuclear structures that contain components that serve to prepare piRNA precursors for loading onto Piwi (Olivieri et al. 2010; Saito et al. 2010b; Hirakata et al. 2019; Qi et al. 2011; Szakmary et al. 2009). Yb is the central Yb body-forming component that also forms granules in S2R+ cells (**Figure 4K**). Previously, a binding hierarchy for Yb body assembly in ovarian somatic cells (OSCs) was proposed, in which Yb binds to Armi and Armi recruits a complex composed of Vret and SoYb (Hirakata et al. 2019). While we observed individual Yb - Armi and Vret - SoYb interactions, our pairwise interaction tests did not detect a link between these complexes. We therefore speculate that additional factors may be required to link the Vret-SoYb complex to Yb-Armi, such as RNA or another Yb body protein component that remains to be identified. Alternatively, a specific conformational state (e.g., RNA-induced) of one or more of the components may be a prerequisite for the assembly of a quaternary complex.

### Possible functional implications of the Vret - BoYb/SoYb complexes

*Drosophila* Vreteno and the BoYb and SoYb proteins have been poorly characterized at the molecular and mechanistic levels. Our interaction studies and structural characterization of the corresponding Vret-BoYb and Vret-SoYb complexes revealed a binding interface, in which the RRM domain of Vret contacts the N-terminal RecA-like domain of the BoYb or SoYb helicase core. Proteins or protein domains that contact the core of DEAD-box RNA helicases can act as modulators of helicase activity, including MIF4G domains (Buchwald et al. 2013; Oberer et al. 2005; Hilbert et al. 2011; Andreou and Klostermeier 2014; Schütz et al. 2008), eLOTUS domains (Jeske et al. 2017), and RRM domains (Samatanga et al. 2017; Wang et al. 2006; Rudolph and Klostermeier 2009; Wurm et al. 2021). Although no *in vitro* activity of BoYb or SoYb has so far been demonstrated, it is tempting to speculate that the RRM domain of Vret might contribute to RNA binding or even modulate the putative enzymatic activities of BoYb or SoYb. Interestingly, all RRM domains previously known to modify the activity of their bound helicase bind to a different helicase surface than that observed in the Vret - BoYb/SoYb complex (**Supplementary Figure 8A**).

Our studies showed that the C-terminal part of Vret contacts the CS domain of BoYb or SoYb. CS domains were originally described in Hsp90 co-chaperones (Garcia-Ranea et al. 2002). They can bind to the closed ATP-bound or the open nucleotide-free form of the N-terminal domain of Hsp90. For example, the co-chaperone p23 (Sba1 in yeast) binds to the closed, ATP-bound form of Hsp90 and stabilizes this state (Ali et al. 2006; Sullivan et al. 1997; Kosano et al. 1998; Dittmar et al. 1997), whereas the CS domain of the co-chaperone Sgt binds the open form (Zhang et al. 2008). Based on a DALI search (Holm and Rosenström 2010), the predicted structures of the CS domains of BoYb and SoYb are most similar to the structure of p23. Superimposing the BoYb-CS - Vret-C complex on the crystal structures of the p23-Hsp90 or the Sgt-CS - Hsp90 complexes revealed that Vret binds to a surface of the CS domain that is opposite to the corresponding Hsp90-binding regions of p23 or Sgt (**Supplementary Figure 8B**). Whether the CS domains of BoYb and/or SoYb actually bind to Hsp83 in any conformational state remains to be tested. Nevertheless, it is tempting to speculate that the CS domains of BoYb and SoYb not only bind to Vreteno, but may also function in Hsp90 recruitment. This is particularly exciting, since Hsp83 (the *Drosophila* ortholog of Hsp90) and the cochaperone Shutdown (FKBP6 in mouse) have been identified as critical components of the piRNA pathway (Specchia et al. 2010; Preall et al. 2012; Olivieri et al. 2012; Xiol et al. 2012). Since the Hsp90 chaperone machinery is essential for loading miRNAs or siRNAs onto Ago proteins (Iwasaki et al. 2010, 2015; Miyoshi et al. 2010; Iki et al. 2010; Naruse et al. 2018), it has been proposed that this machinery plays a similar role in loading piRNAs onto PIWI proteins (Czech and Hannon 2016; Olivieri et al. 2012). Based on previous genetic data, the SoYb-Vret complex is potentially important for the function of somatic piRNA biogenesis and the BoYb-Vret complex for germline function (Handler et al. 2011). Thus, the function of BoYb and SoYb may be linked to PIWI protein loading in somatic and germline cells, an exciting hypothesis that remains to be tested in the future.

Although the sequences and domain organization of BoYb and SoYb are similar to the Yb protein (**compare Figure 2A with Figure 4J**), we did not detect any interaction of Yb with Vret. The lack of binding may be explained by the structural insights gained from the Vret-BoYb/SoYb complexes. First, Yb lacks a CS domain, eliminating one of the Vret-BoYb/SoYb interfaces. Second, the surface of the N-terminal RecA-like domain of Yb, which contacts the RRM of Vret in BoYb/SoYb, is less hydrophobic compared to that on BoYb/SoYb and instead positively charged (**Supplementary Figure 9A**), possibly preventing complex formation. Notably, AlphaFold-Multimer analysis did not result in a high confidence structure prediction of the Yb-Vret complex, as shown by the PAE plots (**Supplementary Figure 9B**). Taken together, our analysis may explain why Vret binds to the Yb-related proteins BoYb and SoYb, but not to Yb. Although Yb, BoYb and SoYb share a similar domain organization and extensive sequence similarity, the specific binding preference of Vret for BoYb and SoYb but not for Yb and the Yb-specific binding of Armi may explain the different biological roles of BoYb and SoYb on the one hand and Yb on the other.

### Tejas is a Vasa helicase effector that might serve as central hub in ping-pong amplification

The domain organization of Tejas is very similar to that of Tapas, both containing an eLOTUS domain that binds and stimulates the Vasa ATPase (Jeske et al. 2017), and one or more additional eTudor domains. However, Tapas cannot replace the function of Tejas *in vivo* (Patil et al. 2014). Our identification of Maelstrom and Krimper as Tejas, but not Tapas, binding partners and the functional consequence of the formation of these protein complexes may therefore explain the distinct *tejas* and *tapas* phenotypes.

In this study, we identified Maelstrom as an additional binding partner of Tejas. Previously, the Mael domain of Maelstrom was shown to be necessary and sufficient for the nuclear function of Maelstrom in OSCs (Sienski et al. 2012; Matsumoto et al. 2015). Tejas localizes to the nuage (Patil and Kai 2010) and is the only Maelstrom interaction partner we detected in our ReLo interaction screen using cytoplasmic and mitochondrial surface proteins (nuclear proteins were not tested). Tejas binds to the TIR at the very C-terminus of Maelstrom (aa 413-459), and therefore disruption of the Tejas-Maelstrom interaction by mutation or deletion of the TIR *in vivo* should not affect the nuclear function of Maelstrom. This may allow the uncoupling of the nuclear from a cytoplasmic function of Maelstrom in in vivo studies. The *Bombyx* Maelstrom ortholog localizes to the cytoplasm, links the proteins Spindle E and the *Bombyx* Aub ortholog Siwi, and is required for the formation of piRNA-loaded Ago3 (Namba et al. 2022). Compared to the *Drosophila* protein, the C-terminal region of *Bombyx* Maelstrom is shorter and lacks a potential Tejas-interacting region (**Supplementary Figure 7I**). In addition, we did not detect any interaction between *Drosophila* Maelstrom and Spindle E or Aubergine in our PPI screen, suggesting that the interactions may either require RNA, a specific conformational state, or that the molecular mechanisms underlying the function of Maelstrom are likely different between *Bombyx* and *Drosophila*. Future *in vivo* analysis of a Tejas-binding mutant of *Drosophila* Maelstrom is likely to provide new insights into the cytoplasmic function of Maelstrom.

Krimper is another previously unknown Tejas binding partner that we identified in our PPI screen. Similar to Tejas, Krimper is a germline-specific TDRD that is concentrated in the perinuclear nuage of nurse cells (Lim and Kai 2007). Krimper has been shown to bind to the PIWI proteins Aub and Ago3 and to coordinate Aub/Ago3-driven ping-pong amplification of piRNAs by preventing homotypic Aub-Aub ping-pong (Sato et al. 2015; Webster et al. 2015). Here, we confirmed the sDMA-dependent interaction between Krimper and Aub and the sDMA-independent interaction between Krimper and Ago3 (**Figure 1**). In addition, we found that Tejas can simultaneously bind to Krimper, Spindle E and the previously known Tejas binder Vasa (**Figure 7**). We previously discovered that it is the eLOTUS domain of Tejas that binds to Vasa and stimulates the Vasa ATPase activity in vitro (Jeske et al. 2017). *Drosophila* Vasa has previously been functionally linked to Aub (Xiol et al. 2014), and the *Bombyx* Vasa ortholog was proposed to be required for the release of the cleaved target RNA strand from piRNA-loaded Siwi (Nishida et al. 2015). In our ReLo screen, we did not detect any interaction between Vasa and Aub. Instead, we discovered that the Tejas - Krimper complex may provide a direct link between Vasa and Aub. Together with our and recent data (Lin et al. 2023) on the Tejas-Spindle E interaction, we conclude that Tejas is a critical core component of nuage.

### Functional insights into non-canonical eTudor domains

Many TDRDs involved in the piRNA pathway contain eTudor domains that lack an intact aromatic cage (**Supplementary Figure 1C**) (Handler et al. 2011) and are therefore unlikely to bind to their partner in an sDMA-dependent, canonical manner. The functions of these non-canonical eTudor domains in *Drosophila* remain unknown. In our screen, we detected interaction partners for several non-canonical eTudor domains, and through structural analysis, we discovered that they use several unusual surfaces to interact, all of which are distinct from the canonical peptide-binding cleft formed by the Tudor and the SN-fold in eTudor domains with an intact aromatic cage (**Supplementary Figure 10**). In the complex between Vreteno and BoYb or SoYb, an α-helix connecting the Tudor domain to the SN-fold in the eTudor domain of Vreteno is mainly involved in the interaction with the CS domain of BoYb or SoYb, whereas the eTudor domain of Tejas binds to Maelstrom or Krimper through an α-helix located laterally of the SN-like fold. In addition, the Tejas eTudor domain is extended N-terminally by a short, conserved stretch that binds to Spindle E. We have shown that Tejas can bridge the interaction between Spindle E and Maelstrom or Krimper, suggesting that the Tejas eTudor domain can accommodate more than one interaction partner. Recent work in *C. elegans* has revealed two additional non-canonical eTudor domain interactions that use yet a different eTudor domain surface to interact (Podvalnaya et al. 2023; Bronkhorst et al. 2023) (**Supplementary Figure 10**). In summary, we conclude that eTudor domains lacking an intact aromatic cage use a variety of binding modes to interact with binding partners.

## ACKNOWLEDGEMENTS

We thank Sarah Barnes, Eva Boberlin, Melina Dabek, Yoan Genchev, Carolin Höfer, Jana Kubíková, Levi Miederer, Marius Müldner, Katharina Müller, Sandra Müller, Corinna Nowak, Rebecca Reinig, and Xiaohan Zhao for technical assistance provided with some of the experiments and data analyses. We thank the Nikon Imaging Center of the University of Heidelberg for access to microscopes. We thank the data storage service SDS@hd, supported by the Ministry of Science, Research and the Arts Baden-Württemberg (MWK) and the German Research Foundation (DFG) through the grants INST 35/1314-1 FUGG and INST 35/1503-1 FUGG. This work was funded by the Emmy Noether Program of the German Research Foundation (DFG; JE-827/1-1 to M.J.).

## METHODS

### DNA constructs

The vectors for ReLo assays have been described previously (Jeske et al. 2017; Behm-Ansmant et al. 2006; Kubíková et al. 2023; Salgania et al. 2024). Detailed information on all plasmids used in this study is provided in **Supplementary Table 1**.

### Protein-protein interaction assays

ReLo assays have been described previously (Salgania et al. 2024). The single cell(s) shown in the figures are representative of the entire co-transfected cell population. Protein expression, purification and GST pull-down assays were performed in 100 µl reactions as described previously (Kubíková et al. 2023). For GST fusion constructs, 1 nmol and for MBP fusion constructs, 2 nmol (Figure 6D) and 5 nmol (Figure 6F), respectively, were used.

### Structure prediction with AlphaFold

The ColabFold v1.5.2 web interface (Mirdita et al. 2022) was used for structure prediction with default settings except for model_type, which was changed from “auto” to “alphaFold_multimer_v2” (earlier searches) or “alphaFold_multimer_v3” (later searches). Structures were visualized with PyMol. The model of the Armi-Daed complex bound to dsRNA was generated with AlphaFold 3 (Abramson et al. 2024) using the web interface with standard settings.

### Declaration of generative AI and AI-assisted technologies in the writing process

During the preparation of this work the authors used DeepL (deepl.com) in order to review the English language of the text. After using this tool, the authors reviewed and edited the content as needed and take full responsibility for the content of the publication.

