## Supplementary Data for "Molecular insight into the *Drosophila* piRNA pathway network through a combination of systematic protein interaction screening and structural prediction"

[2] Present address: Groningen Biomolecular Sciences and Biotechnology Institute, University of Groningen, 9747 AG Groningen, The Netherlands

*Correspondence

Inventory:

Supplementary Figure 1
Supplementary Figure 2 (6 pages)
Supplementary Figure 3
Supplementary Figure 4
Supplementary Figure 5
Supplementary Figure 6
Supplementary Figure 7
Supplementary Figure 8
Supplementary Figure 9
Supplementary Figure 10

Supplementary Table 1

Supplementary References

### SUPPLEMENTARY FIGURES

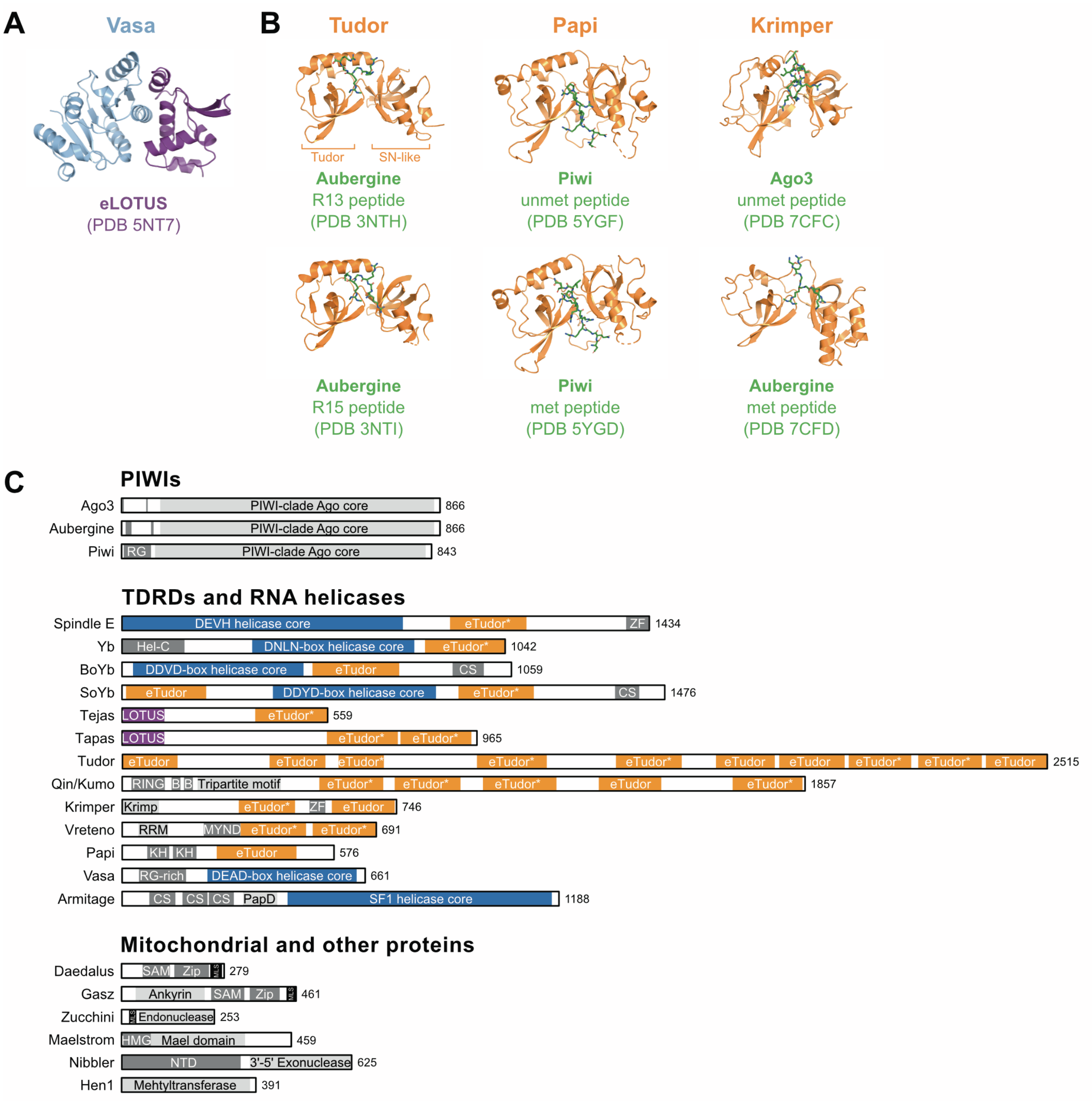

Supplementary Figure 1. **Experimental structural information on *Drosophila* piRNA pathway protein complexes operating in the cytoplasm.**
(A) Complex formed by an eLOTUS domain and the C-terminal RecA-like domain of Vasa.
(B) Complexes formed by eTudor domains and bound peptides.
(C) Domain organization of the cytoplasmic and mitochondrial surface proteins used in this study. eTudor domains lacking an intact aromatic cage are indicated with an asterisk (*).

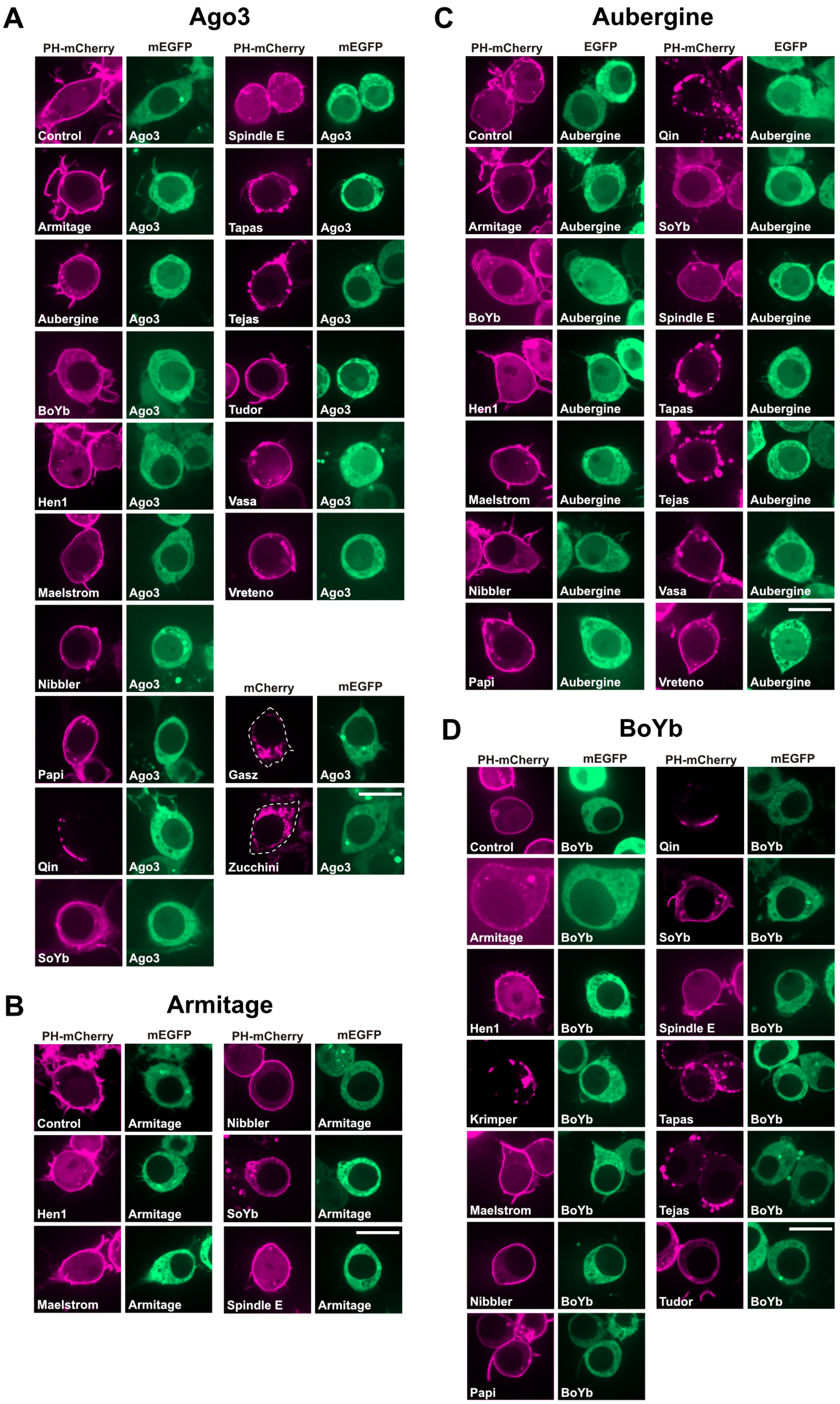

Supplementary Figure 2. **ReLo assay-based PPI screen of cytoplasmic and surface mitochondrial piRNA pathway factors**. (part 1; legend on page 8)

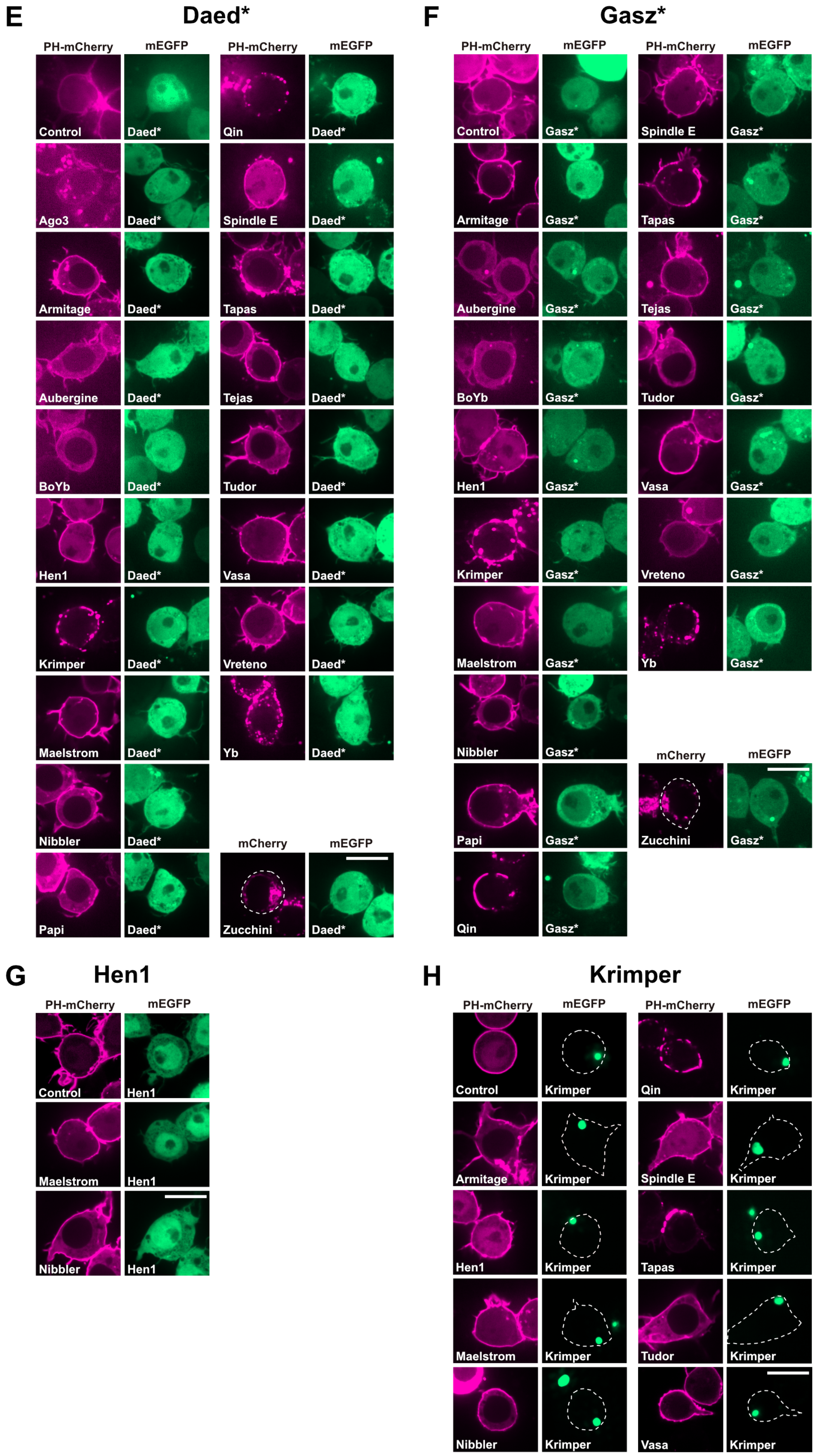

Supplementary Figure 2. **ReLo assay-based PPI screen of cytoplasmic and surface mitochondrial piRNA pathway factors**. (part 2; legend on page 8)

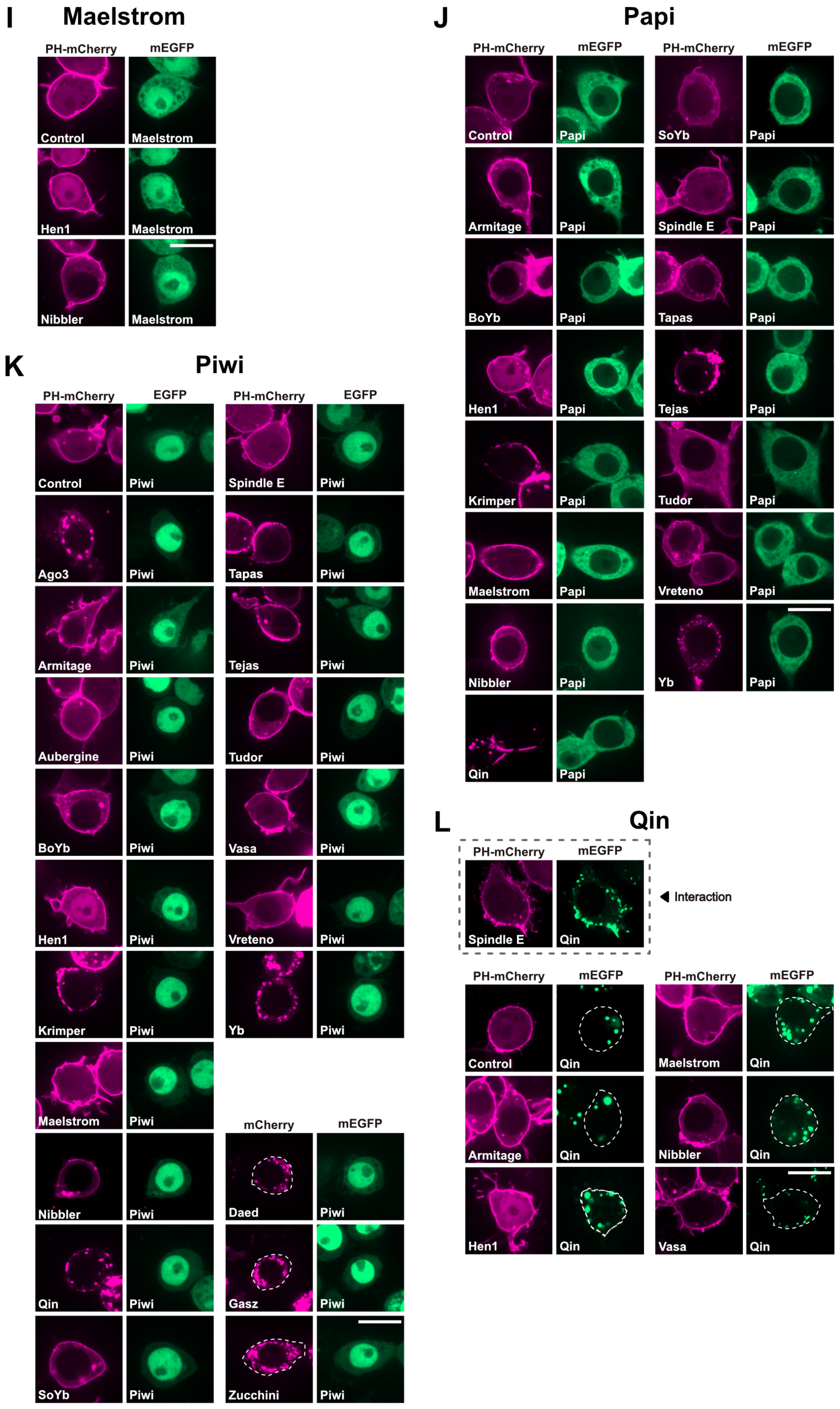

Supplementary Figure 2. **ReLo assay-based PPI screen of cytoplasmic and surface mitochondrial piRNA pathway factors**. (part 3; legend on page 8)

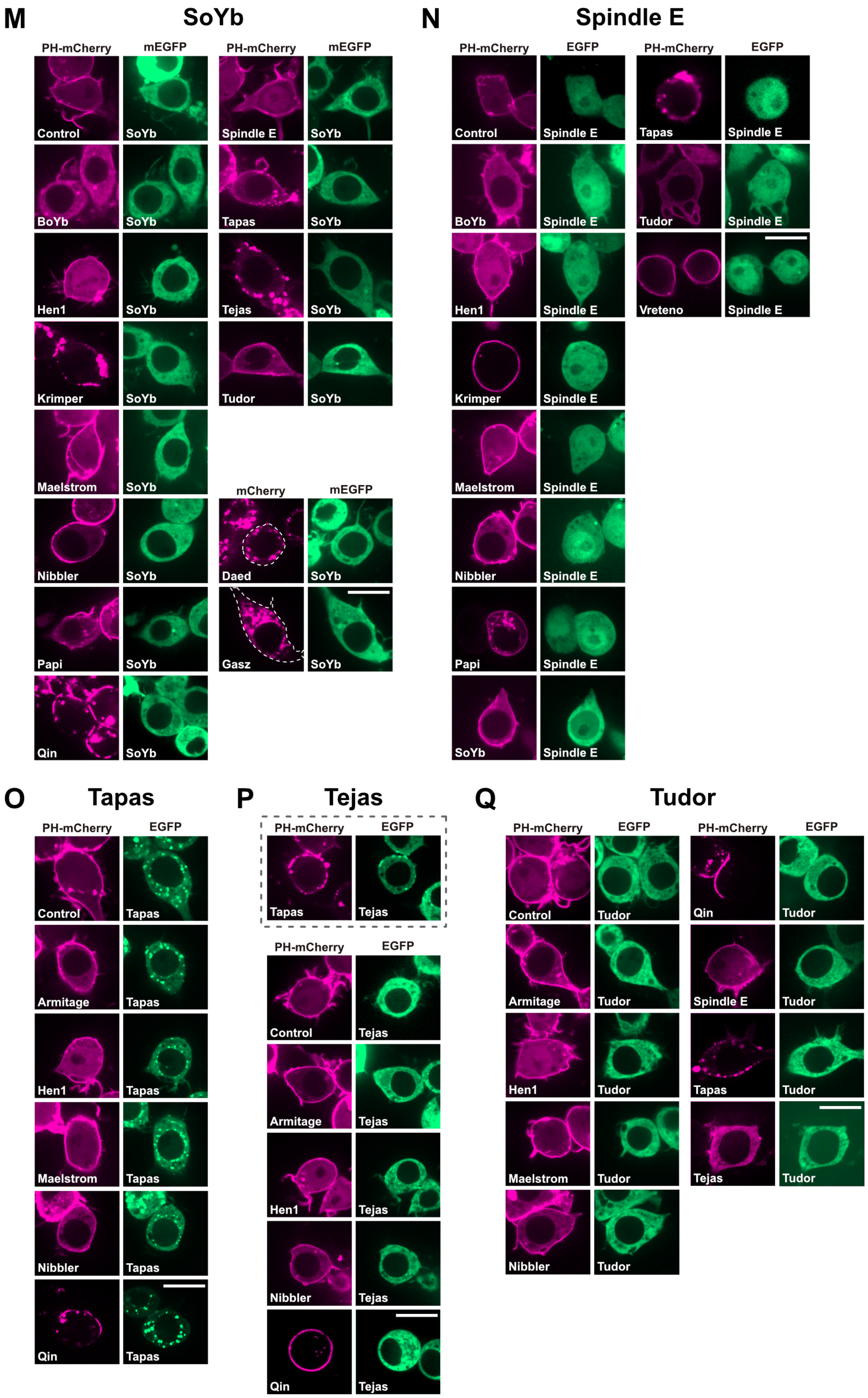

Supplementary Figure 2. **ReLo assay-based PPI screen of cytoplasmic and surface mitochondrial piRNA pathway factors**. (part 4; legend on page 8)

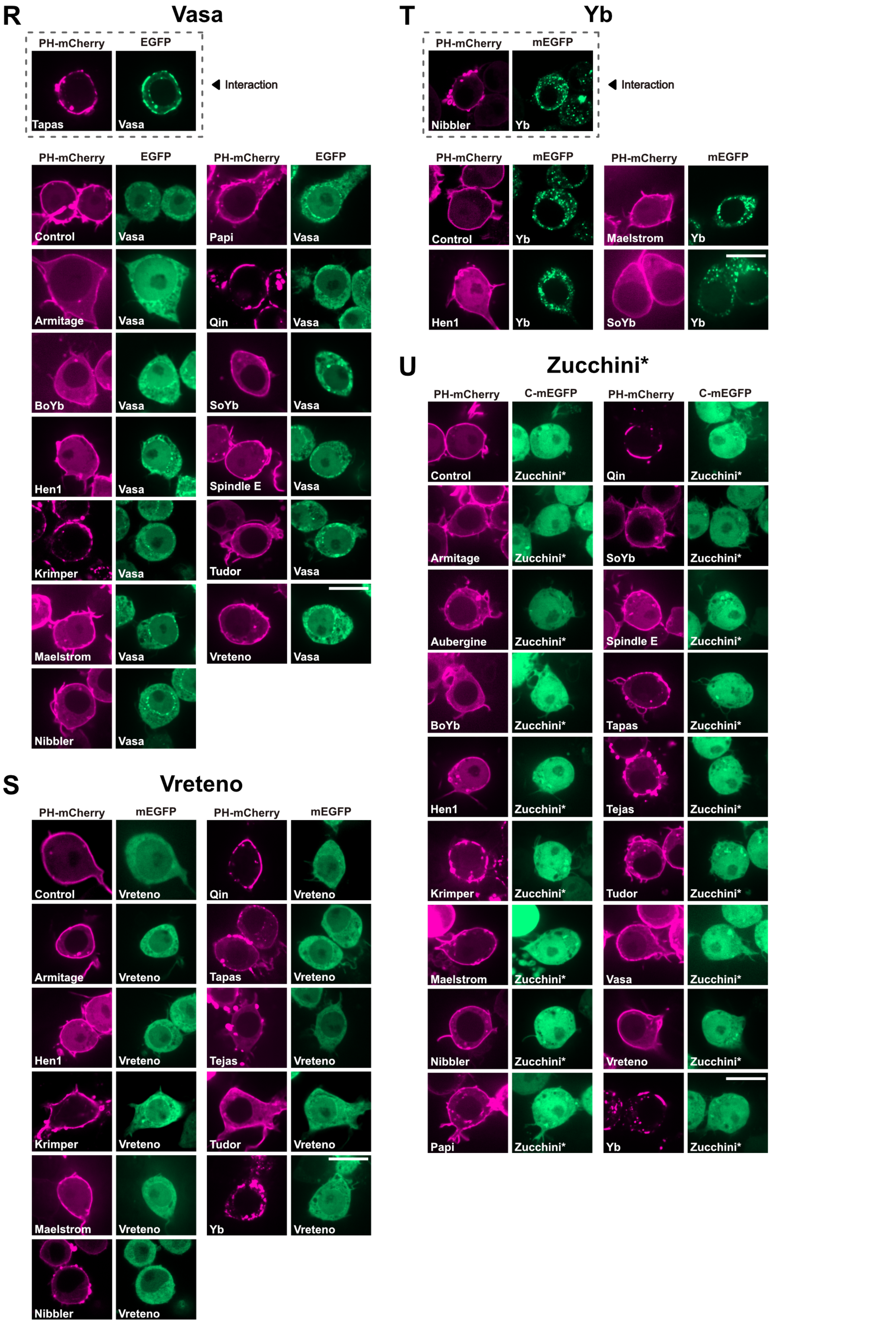

Supplementary Figure 2. **ReLo assay-based PPI screen of cytoplasmic and surface mitochondrial piRNA pathway factors**. (part 5; legend on page 8)

**V** **Controls**

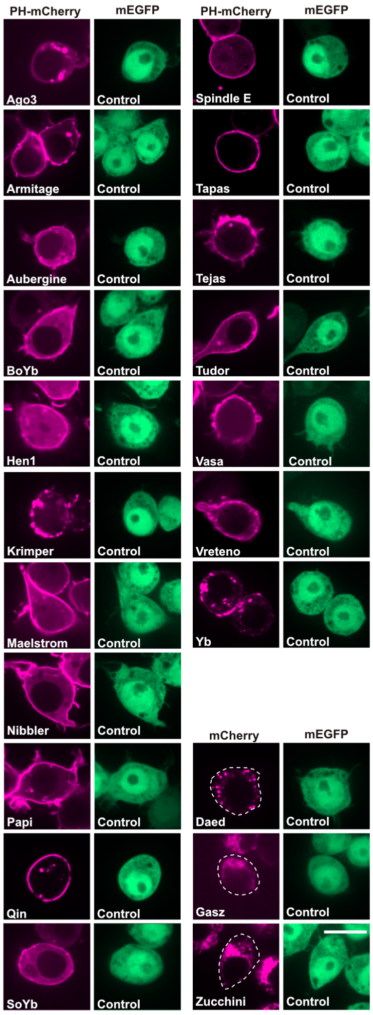

Supplementary Figure 2. **ReLo assay-based PPI screen of cytoplasmic and mitochondrial surface piRNA pathway factors**. (part 6)
Relocalization to the proteins indicated in the images was tested for the following proteins (in alphabetical order): Ago3 (A), Aubergine (B), Armitage (C), BoYb (D), Daed (E), Gasz (F), Hen1 (G), Krimper (H), Maelstrom (I), Papi (J), Piwi (K), Qin (L), SoYb (M), Spindle E (N), Tapas (O), Tejas (P), Tudor (Q), Vasa (R), Yb (S), Vreteno (T), Zucchini (U). (V) Control transfections are shown. Note that reciprocal tagging was not tested in all cases. The asterisk (*) indicates when the mitochondrial localization signal was deleted from the protein. Scale bars are 10 μm.

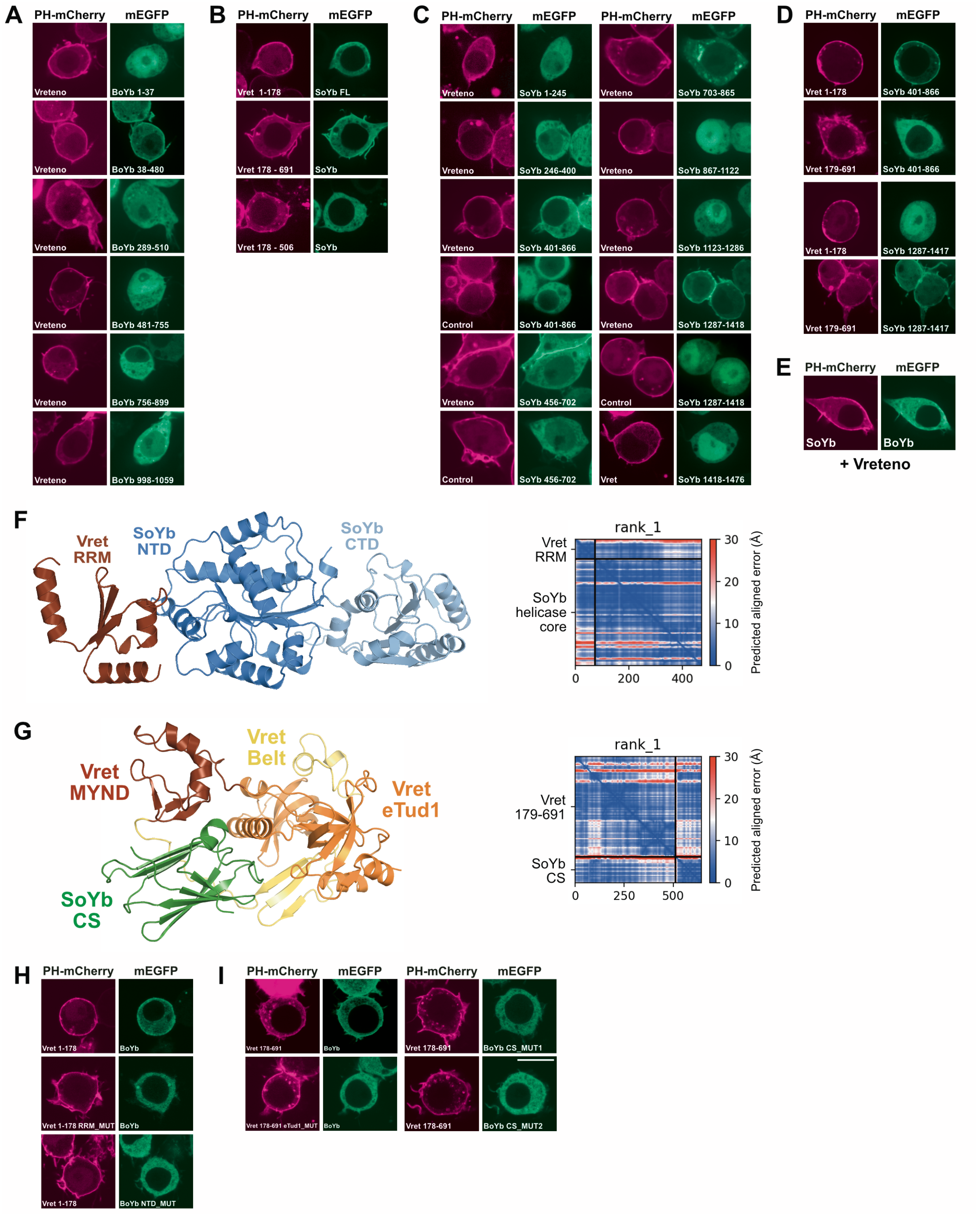

Supplementary Figure 3. **The interactions between Vreteno and BoYb or SoYb.** (see next page)

Supplementary Figure 3. **The interactions between Vreteno and BoYb or SoYb**.
(A) Additional data related to the Vret-BoYb interaction mapping. Controls are shown in **Figure 2**.
(B) Both the N- and C-terminal portions of Vret are required for SoYb binding.
(C) Mapping data showing that the N-terminal RecA-like domain (aa 401-702) and the CS domain (aa 1287-1418) of SoYb contribute to Vret binding.
(D) The N-terminal part of Vret binds the NTD of SoYb and the C-terminal part of Vret binds the CS domain of SoYb.
(E) Vret did not bind to BoYb and SoYb simultaneously.
(F) AlphaFold-Multimer-generated structural model of the complex consiting of the Vret-RRM (aa 43-117) and the helicase core of SoYb (aa 460-856). The PAE plot indicates the high confidence of the model.
(G) AlphaFold-Multimer-generated structural model of the complex composed of the C-terminal part of Vret (aa 179-691) and the CS domain of SoYb (aa 1287-1417). The PAE plot indicates the high confidence of the model.
(H) Mutations that abolished the interaction between the RRM of Vret and the NTD of BoYb.
(I) Mutations that abolished the interaction between the C-terminal part of Vret and the CS domain of BoYb.
Scale bars are 10 μm.

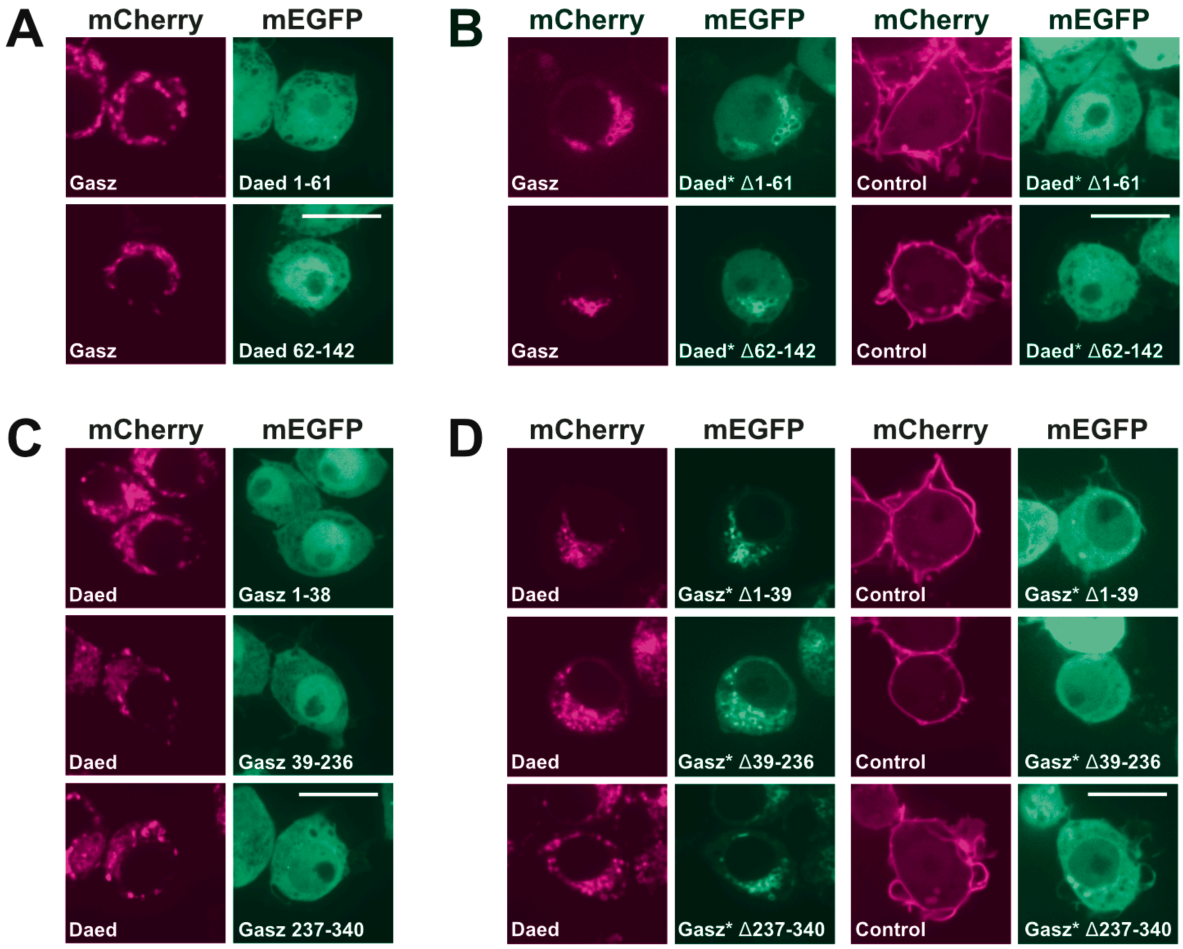

Supplementary Figure 4. **The interaction between Gasz and Daed**.
(A-D) Additional data related to the mapping of the Gasz-Daed complex. The asterisk (*) indicates when the mitochondrial localization signal was deleted from the protein. Scale bars are 10 μm.

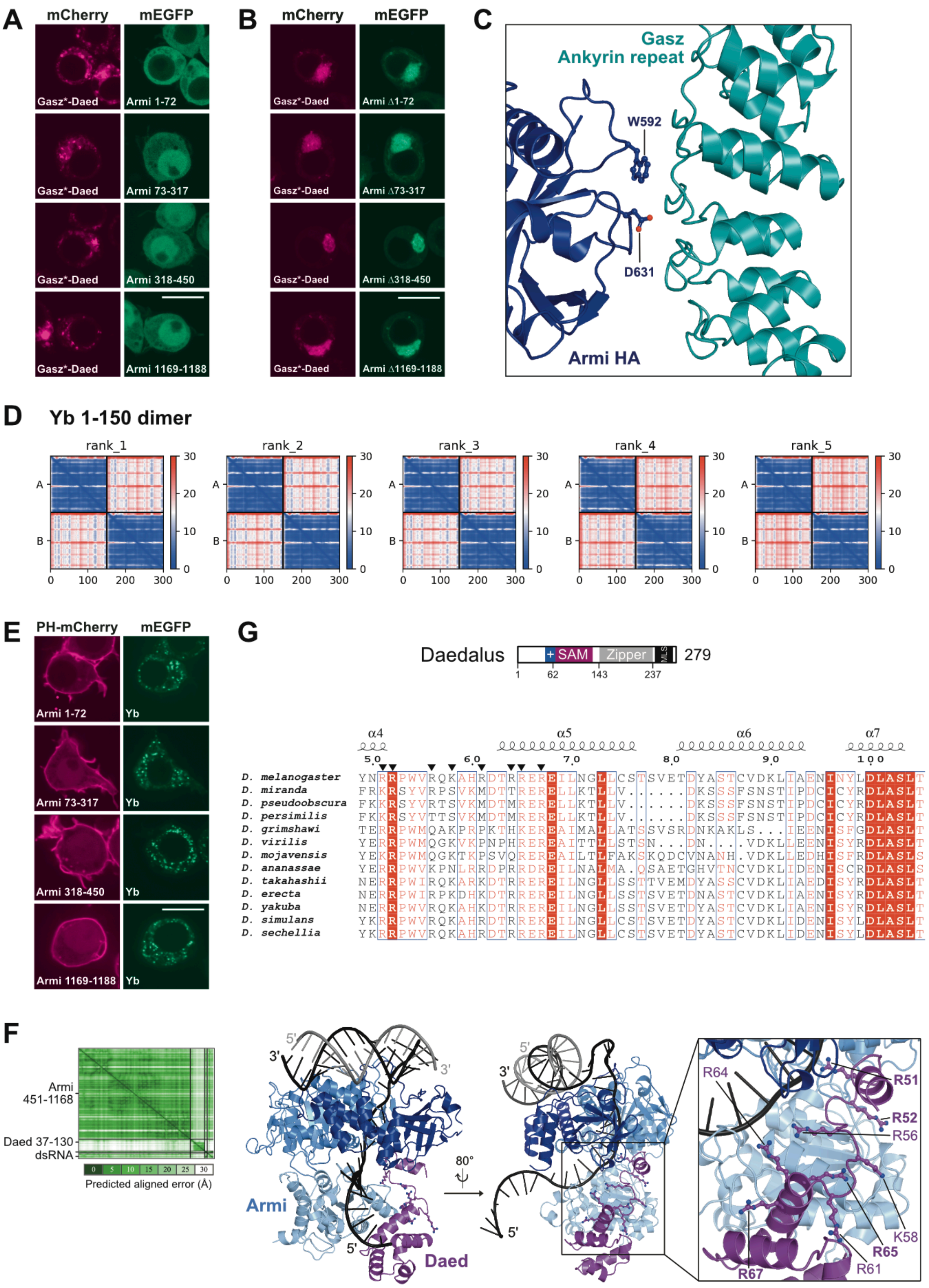

Supplementary Figure 5. **The Gasz-Daed-Armi complex** (legend on next page).

Supplementary Figure 5. **The Gasz-Daed-Armi complex** (continued).
(A, B) Additional data related to the mapping of the Armi regions that bind to the Gasz-Daed complex. The asterisk (*) indicates when the mitochondrial localization signal was deleted from the protein.
(C) Detail of the predicted interface formed by the HA domain of Armi and the Ankyrin domain of Gasz. Residues mutated for the subsequent studies shown in **Figure 4I** are highlighted in ball-and-stick representation.
(D) PAE plots showing that a high confidence model of a Yb 1-150 dimer was not obtained.
(E) Additional data related to the mapping of the Armi regions that bind to Yb.
Scale bars are 10 μm.
(F) AlphaFold 3-generated structural model of the complex formed by the Armi helicase core (aa 451-1168), the N-terminally extended SAM domain of Daed (aa 37-130), and a 5'-overhang dsRNA oligo as previously used in unwinding assays (Ishizu et al. 2019). Positively charged residues that point or are close to the nucleic acid are highlighted in ball-and-stick representation. Chemically conserved residues are shown in bold.
(G) Multiple sequence alignment showing the positively charged region extending N-terminally from the SAM domain of Daed. Charged residues that point to the site where nucleic acid is potentially bound by Armi and Daed are indicated by a black arrowhead at the top of the alignment.

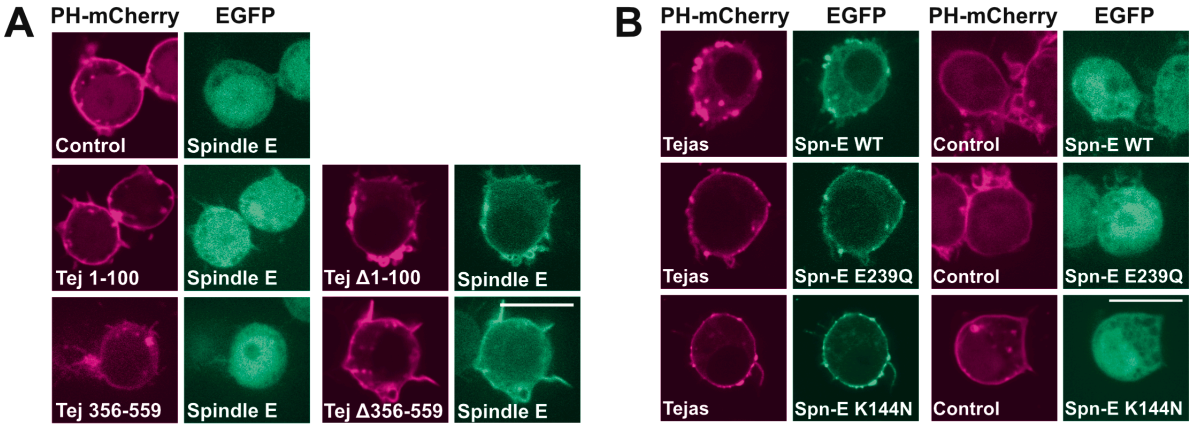

Supplementary Figure 6. **The interaction between Tejas and Spindle E**.
(A) Additional data related to the mapping of the Tejas-Spindle E interaction.
(B) Mutations on the Spindle E surface that were not sufficient to abolish the interaction with Tejas.

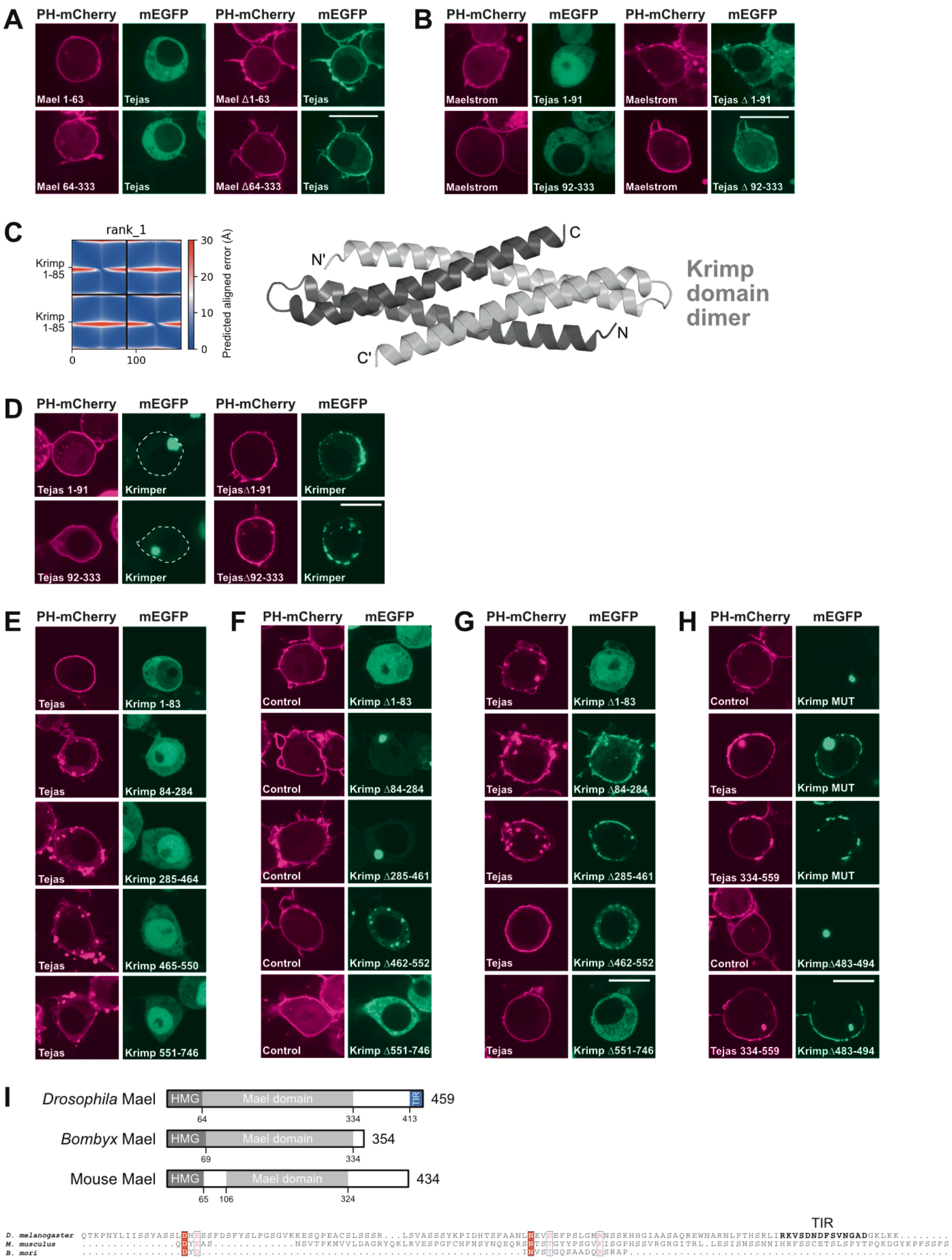

Supplementary Figure 7. **The interaction between Tejas and Maelstrom or Krimper**. (next page)

Supplementary Figure 7. **The interaction between Tejas and Maelstrom or Krimper**.
(A, B) Additional data related to the mapping of the Tejas-Maelstrom complex.
(C) AlphaFold-Multimer-generated structural model of the Krimp domain dimer. The PAE plot indicates the high confidence of the model.
(D) Additional data related to the mapping of the Tejas-Krimper complex.
(E) None of the Krimper fragments showed any interaction with Tejas.
(F) Subcellular localization of the Krimper deletion constructs.
(G) The interaction with Tejas seems to require the Krimp domain (aa 1-83) and the C-terminal part (aa 462-746) of Krimper.
(H) The Krimper mutations and deletions of the Tejas-interacting region predicted by AlphaFold-multimer are not sufficient to abolish the Tejas interaction, suggesting the contribution of additional regions in Krimper.
(I) Domain organization of Maelstrom from *Drosophila*, *Bombyx* and mouse. Multiple sequence alignment (Edgar 2004) of the C-terminal unstructured regions of Maelstrom.
Scale bars are 10 μm.

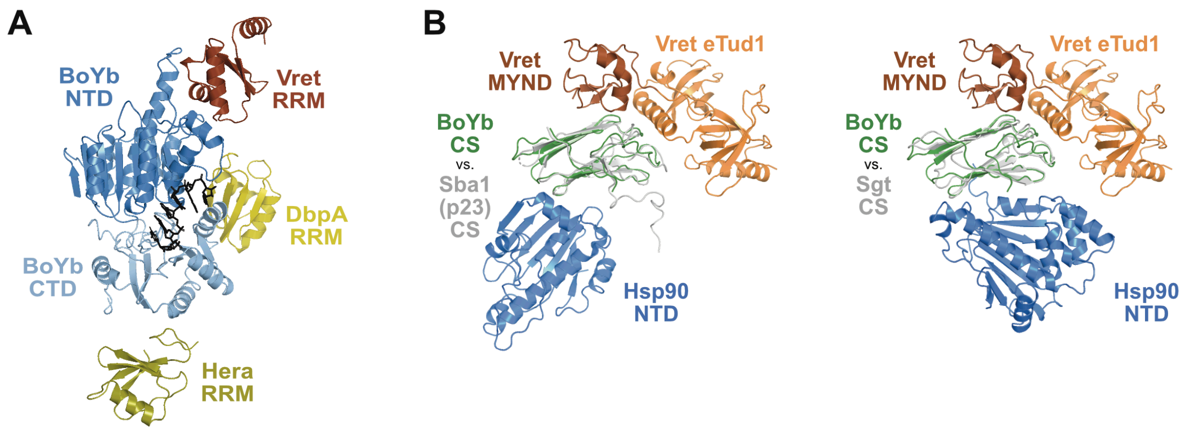

Supplementary Figure 8. **Analysis of the predicted structures of the BoYb-Vret complex.**

(A) Composite structural model visualizing the binding positions of the RRM of Vret to the BoYb-NTD, the RRM of DbpA at the DbpA core, and the RRM of Hera to the Hera-CTD. The BoYb helicase closed core was generated by superimposing the BoYb-NTD - Vret complex generated by AlphaFold-Multimer on the BoYb closed core modeled with SWISS-MODEL (Waterhouse et al. 2018) using the Vasa closed core (PBD 2DB3) as a template. The DbpA and Hera structures (PDBs 7PLI and 3I32, respectively) were superimposed on the BoYb helicase core and their RRMs were extracted. The RNA oligo was extracted from PDB 7PLI.

(B) Superimposition of the AlphaFold-Multimer generated BoYb-CS - Vret-CTD complex with the yeast Sba1-CS - Hsp90-NTD complex (PDB 2CG9; left panel) or the *Arabidopsis* Sgt-CS - Hsp90-NTD complex (PBD 2XCM, right panel) structures. The models were superimposed on the basis of the CS domains.

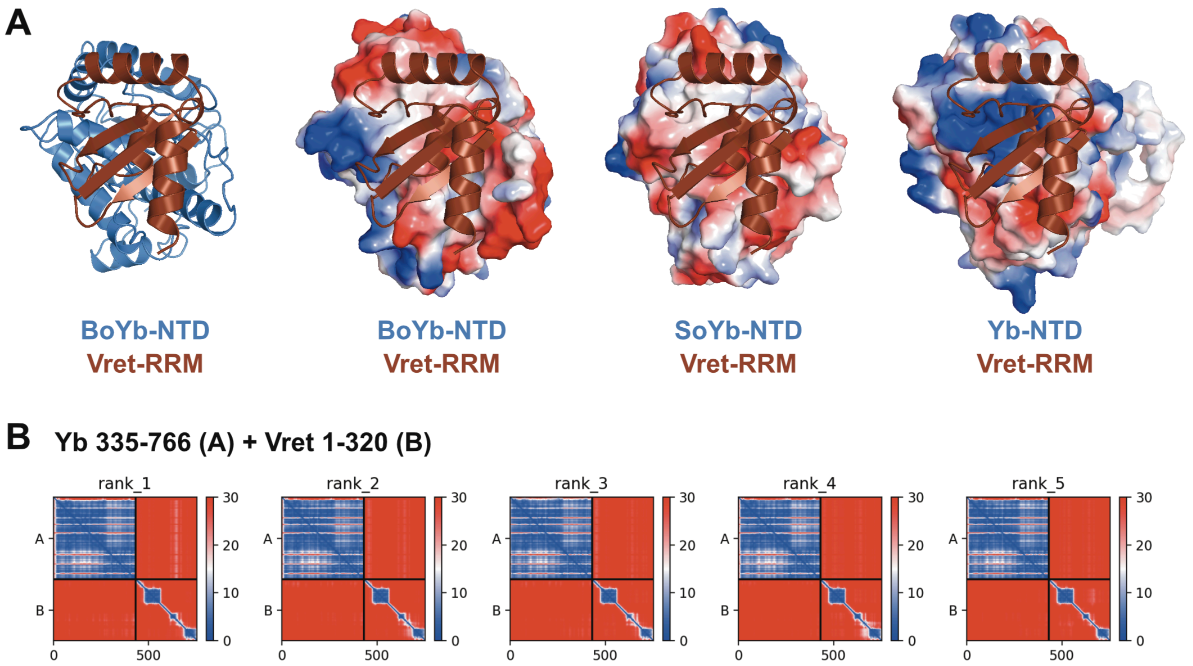

Supplementary Figure 9. **Structural comparison of BoYb and SoYb with Yb.**
(A) Comparison of the charge of the BoYb and SoYb surfaces involved in the Vret interaction.
(B) Predicted aligned error plots for the structural model prediction run using the Yb helicase core and the N-terminal part of Vret.

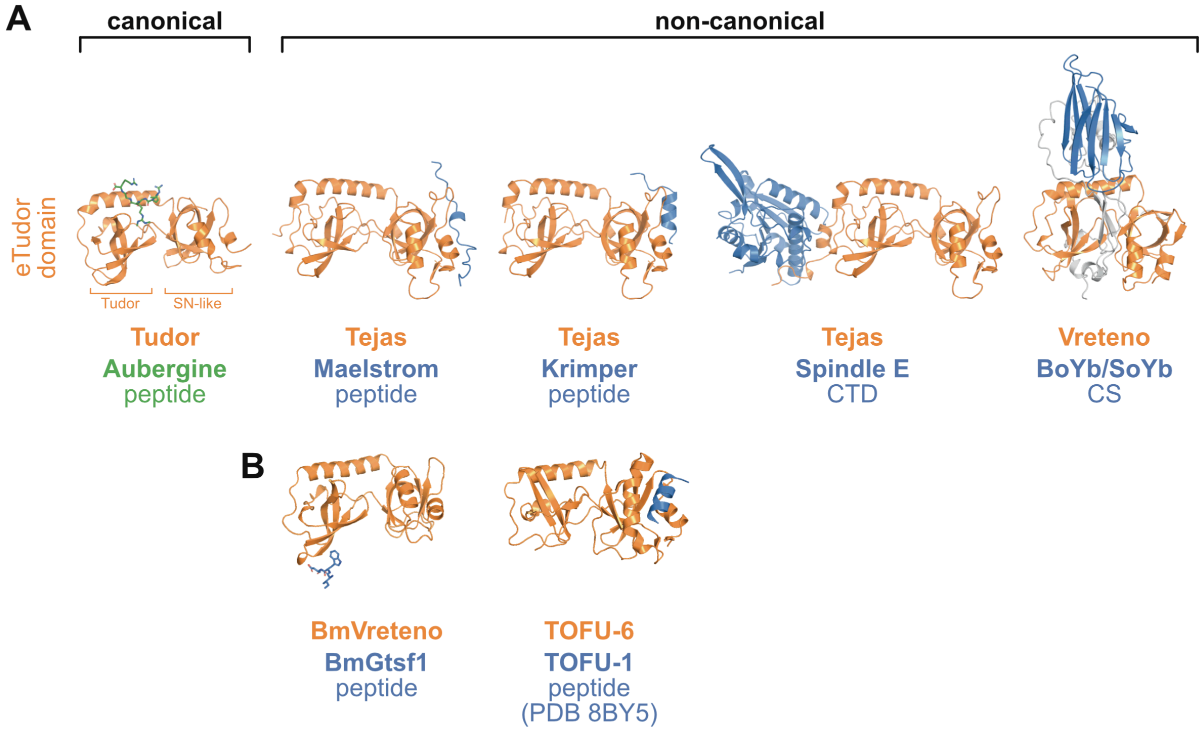

Supplementary Figure 10. **Non-canonical eTudor domains.**
(A) Comparison of non-canonical eTudor domains lacking an intact aromatic cage bound to their interaction partners.
(B) Previously reported non-canonical eTudor domains lacking an intact aromatic cage bound to their interaction partners (Bronkhorst et al. 2023; Podvalnaya et al. 2023).

**Supplementary Table 1. DNA constructs used in this study.**
All constructs are *Drosophila melanogaster* sequences, if not indicated otherwise. The specific protein isoforms (iso) used are indicated.

| **Vector** (insertion site) (code) | **Final DNA construct** | **DNA template information** | **Code** |
|---|---|---|---|
| **pAc5.1-EGFP** (EcoRV) (T5-MJ) | pAc5.1-EGFP-**Aubergine** iso A | (Salgania et al. 2024) | F20-MJ |
| | pAc5.1-EGFP-**Aubergine 4R→K (**R11K/R13K/R15K/R17K) | (Salgania et al. 2024) | HK121 |
| | pAc5.1-EGFP-**Piwi** iso A | *Drosophila* ovarian cDNA | F30-MJ |
| | pAc5.1-EGFP-**Piwi Δ1-90** | Sequence specific deletion in pAc5.1-EGFP-Piwi | HK121 |
| | pAc5.1-EGFP-**Spindle E** iso A | *Drosophila* ovarian cDNA | F40-MJ |
| | pAc5.1-EGFP-**Spindle E 1-765** | pAc5.1-EGFP-Spindle E | RR138 |
| | pAc5.1-EGFP-**Spindle E 766-1434** | pAc5.1-EGFP-Spindle E | RR139 |
| | pAc5.1-EGFP-**Spindle E E239Q** | Site directed mutagenesis of pAc5.1-EGFP-Spindle E | RR143 |
| | pAc5.1-EGFP-**Spindle E K144N** | Site directed mutagenesis of pAc5.1-EGFP-Spindle E | RR144 |
| | pAc5.1-EGFP-**Tapas** iso D | (Jeske et al. 2017) | F34-MJ |
| | pAc5.1-EGFP-**Tejas** iso A | (Jeske et al. 2017) | F33-MJ |
| | pAc5.1-EGFP-**Tejas 1-91** | pAc5.1-EGFP-Tejas | EL78 |
| | pAc5.1-EGFP-**Tejas 92-333** | pAc5.1-EGFP-Tejas | EL81 |
| | pAc5.1-EGFP-**Tejas 334-559** | pAc5.1-EGFP-Tejas | EL80 |
| | pAc5.1-EGFP-**Tejas Δ1-91** | Sequence specific deletion in pAc5.1-EGFP-Tejas | EL85 |
| | pAc5.1-EGFP-**Tejas Δ92-333** | Sequence specific deletion in pAc5.1-EGFP-Tejas | EL86 |
| | pAc5.1-EGFP-**Tejas Δ334-559** | Sequence specific deletion in pAc5.1-EGFP-Tejas | EL87 |
| | pAc5.1-EGFP-**Tejas D320K** | Site directed mutagenesis of pAc5.1-EGFP-Tejas | RR218 |
| | pAc5.1-EGFP-**Tejas A332E/V333E** | Site directed mutagenesis of pAc5.1-EGFP-Tejas | RR243 |

| | | | |
|---|---|---|---|
| | pAc5.1-EGFP-**Tejas V347E** | Site directed mutagenesis of pAc5.1-EGFP-Tejas | RR215 |
| | pAc5.1-EGFP-**Tejas MUT** (V530E/V532E) | Site directed mutagenesis of pAc5.1-EGFP-Tejas | JM260 |
| | pAc5.1-EGFP-**Tudor** iso A | *Drosophila* ovarian cDNA | F38-MJ |
| | pAc5.1-EGFP-**Vasa** iso A | (Salgania et al. 2024) | F15-MJ |
| | pAc5.1-EGFP-**Vasa open** (K295N) | (Salgania et al. 2024) | RR228 |
| **pAc5.1-mEGFP** (EcoRV) (T6-MJ) | pAc5.1-mEGFP-**Ago3** iso D | *Drosophila* ovarian cDNA | JM60 |
| | pAc5.1-mEGFP-**Ago3 Δ1-100** | Sequence specific deletion in pAc5.1-mEGFP-Ago3 | HK144 |
| | pAc5.1-mEGFP-**Ago3 3R→K** (R68K/R70K/R72K) | Site directed mutagenesis of pAc5.1-mEGFP-Ago3 | HK149 |
| | pAc5.1-mEGFP-**Armitage** iso B | *Drosophila* ovarian cDNA | JM63 |
| | pAc5.1-mEGFP-**Armi 1-72** | pAc5.1-mEGFP-Armitage | HK231 |
| | pAc5.1-mEGFP-**Armi 73-317** | pAc5.1-mEGFP-Armitage | HK232 |
| | pAc5.1-mEGFP-**Armi 318-450** | pAc5.1-mEGFP-Armitage | HK233 |
| | pAc5.1-mEGFP-**Armi 451-1168** | pAc5.1-mEGFP-Armitage | HK234 |
| | pAc5.1-mEGFP-**Armi 1169-1188** | pAc5.1-mEGFP-Armitage | HK235 |
| | pAc5.1-mEGFP-**Armi Δ1-72** | Sequence specific deletion in pAc5.1-mEGFP-Armitage | HK236 |
| | pAc5.1-mEGFP-**Armi Δ73-317** | Sequence specific deletion in pAc5.1-mEGFP-Armitage | HK237 |
| | pAc5.1-mEGFP-**Armi Δ318-450** | Sequence specific deletion in pAc5.1-mEGFP-Armitage | HK238 |
| | pAc5.1-mEGFP-**Armi Δ451-1168** | Sequence specific deletion in pAc5.1-mEGFP-Armitage | HK239 |
| | pAc5.1-mEGFP-**Armi Δ1169-1188** | Sequence specific deletion in pAc5.1-mEGFP-Armitage | HK240 |
| | pAc5.1-mEGFP-**Armi L1030E/M1061E** | Site directed mutagenesis of pAc5.1-mEGFP-Armitage | HK291 |

| | | | |
|---|---|---|---|
| | pAc5.1-mEGFP-**BoYb** iso A | *Drosophila* ovarian cDNA | CB17 |
| | pAc5.1-mEGFP-**BoYb 1-37** | pAc5.1-mEGFP-BoYb | EL66 |
| | pAc5.1-mEGFP-**BoYb 1-288** | pAc5.1-mEGFP-BoYb | HK125 |
| | pAc5.1-mEGFP-**BoYb 38-480** | pAc5.1-mEGFP-BoYb | EL68 |
| | pAc5.1-mEGFP-**BoYb 289-510** | pAc5.1-mEGFP-BoYb | HK126 |
| | pAc5.1-mEGFP-**BoYb 481-755** | pAc5.1-mEGFP-BoYb | EL72 |
| | pAc5.1-mEGFP-**BoYb 756-899** | pAc5.1-mEGFP-BoYb | EL69 |
| | pAc5.1-mEGFP-**BoYb 900-997** | pAc5.1-mEGFP-BoYb | EL73 |
| | pAc5.1-mEGFP-**BoYb 998-1059** | pAc5.1-mEGFP-BoYb | EL71 |
| | pAc5.1-mEGFP-**BoYb NTD_MUT** (L192E/L195E/L218E) | Site directed mutagenesis of pAc5.1-mEGFP-BoYb | MD17 |
| | pAc5.1-mEGFP-**BoYb CS_MUT1** (L979E/L980E/L982E) | Site directed mutagenesis of pAc5.1-mEGFP-BoYb | MD18 |
| | pAc5.1-mEGFP-**BoYb CS_MUT2** (L933E/F938E) | Site directed mutagenesis of pAc5.1-mEGFP-BoYb | MD19 |
| | pAc5.1-mEGFP-**BoYb NTD/CS_MUT1** (L192E/L195E/L218E/L979E/L980E/L982E) | Gibson assembly using pAc5.1-mEGFP-BoYb NTD_MUT and pAc5.1-mEGFP-BoYb CS_MUT1 | DB37 |
| | pAc5.1-mEGFP-**Daed 1-61** | pAc5.1-mEGFP-Daed | KO30 |
| | pAc5.1-mEGFP-**Daed 62-142** | pAc5.1-mEGFP-Daed | KO37 |
| | pAc5.1-mEGFP-**Daed 143-236** | pAc5.1-mEGFP-Daed | KO39 |
| | pAc5.1-mEGFP-**Daed*** (∆237-279; ∆MLS) | Sequence specific deletion in pAc5.1-mEGFP-Daed | HK172 |
| | pAc5.1-mEGFP-**Daed*Δ1-61** | Sequence specific deletion in pAc5.1-mEGFP-Daed* | HK245 |
| | pAc5.1-mEGFP-**Daed*Δ62-142** | Sequence specific deletion in pAc5.1-mEGFP-Daed* | HK246 |
| | pAc5.1-mEGFP-**Daed*Δ143-236** | Sequence specific deletion in pAc5.1-mEGFP-Daed* | HK247 |
| | pAc5.1-mEGFP-**Daed*MUT** (F161E/I205E) | Site directed mutagenesis of pAc5.1-mEGFP-Daed* | KO18 |
| | pAc5.1-mEGFP-**Gasz 1-38** | pAc5.1-mEGFP-Gasz | KO33 |
| | pAc5.1-mEGFP-**Gasz 39-236** | pAc5.1-mEGFP-Gasz | KO32 |
| | pAc5.1-mEGFP-**Gasz 237-340** | pAc5.1-mEGFP-Gasz | KO35 |
| | pAc5.1-mEGFP-**Gasz 341-437** | pAc5.1-mEGFP-Gasz | KO34 |
| | pAc5.1-mEGFP-**Gasz*** (∆438-461; ∆MLS) | Sequence specific deletion in pAc5.1-mEGFP-Gasz | XH37 |
| | pAc5.1-mEGFP-**Gasz*Δ1-39** | Sequence specific deletion in pAc5.1-mEGFP-Gasz* | KO1 |
| | pAc5.1-mEGFP-**Gasz*Δ39-236** | Sequence specific deletion in pAc5.1-mEGFP-Gasz* | KO3 |

| | | | |
|---|---|---|---|
| | pAc5.1-mEGFP-**Gasz*Δ237-340** | Sequence specific deletion in pAc5.1-mEGFP-Gasz* | KO2 |
| | pAc5.1-mEGFP-**Gasz*Δ341-437** | Sequence specific deletion in pAc5.1-mEGFP-Gasz* | KO31 |
| | pAc5.1-mEGFP-**Gasz*MUT** (L359E/Y396E) | Site directed mutagenesis of pAc5.1-mEGFP-Gasz* | KO19 |
| | pAc5.1-mEGFP-**Hen1** iso A | *Drosophila* ovarian cDNA | HK155 |
| | pAc5.1-mEGFP-**Krimper** iso A | *Drosophila* ovarian cDNA | HK113 |
| | pAc5.1-mEGFP-**Krimper 1-83** | pAc5.1-mEGFP-Krimper | HK312 |
| | pAc5.1-mEGFP-**Krimper 84-284** | pAc5.1-mEGFP-Krimper | HK321 |
| | pAc5.1-mEGFP-**Krimper 285-461** | pAc5.1-mEGFP-Krimper | HK313 |
| | pAc5.1-mEGFP-**Krimper 462-552** | pAc5.1-mEGFP-Krimper | HK314 |
| | pAc5.1-mEGFP-**Krimper 551-746** | pAc5.1-mEGFP-Krimper | HK315 |
| | pAc5.1-mEGFP-**KrimperΔ1-83** | Sequence specific deletion in pAc5.1-mEGFP-Krimper | HK282 |
| | pAc5.1-mEGFP-**KrimperΔ84-284** | Sequence specific deletion in pAc5.1-mEGFP-Krimper | SB19 |
| | pAc5.1-mEGFP-**KrimperΔ285-461** | Sequence specific deletion in pAc5.1-mEGFP-Krimper | SB21 |
| | pAc5.1-mEGFP-**KrimperΔ462-552** | Sequence specific deletion in pAc5.1-mEGFP-Krimper | SB23 |
| | pAc5.1-mEGFP-**KrimperΔ483-494** | Sequence specific deletion in pAc5.1-mEGFP-Krimper | JM256 |
| | pAc5.1-mEGFP-**KrimperΔ551-746** | Sequence specific deletion in pAc5.1-mEGFP-Krimper | HK137 |
| | pAc5.1-mEGFP-**Krimper MUT** (I483E/L491E/I494E) | Site directed mutagenesis of pAc5.1-mEGFP-Krimper | JM263 |
| | pAc5.1-mEGFP-**Maelstrom** iso C | *Drosophila* ovarian cDNA | HK148 |
| | pAc5.1-mEGFP-**Papi** iso C | *Drosophila* ovarian cDNA | HK114 |
| | pAc5.1-mEGFP-**Qin** iso B | *Drosophila* ovarian cDNA | HK112 |
| | pAc5.1-mEGFP-**SoYb** iso B | *Drosophila* ovarian cDNA | JM88 |
| | pAc5.1-mEGFP-**SoYb 1-245** | pAc5.1-mEGFP-SoYb | EL95 |
| | pAc5.1-mEGFP-**SoYb 246-400** | pAc5.1-mEGFP-SoYb | EL102 |
| | pAc5.1-mEGFP-**SoYb 401-866** | pAc5.1-mEGFP-SoYb | EL103 |
| | pAc5.1-mEGFP-**SoYb 456-702** | pAc5.1-mEGFP-SoYb | HK127 |
| | pAc5.1-mEGFP-**SoYb 703-865** | pAc5.1-mEGFP-SoYb | CB28 |
| | pAc5.1-mEGFP-**SoYb 867-1122** | pAc5.1-mEGFP-SoYb | EL104 |
| | pAc5.1-mEGFP-**SoYb 1123-1286** | pAc5.1-mEGFP-SoYb | EL105 |

| | | | |
|---|---|---|---|
| | pAc5.1-mEGFP-**SoYb 1287-1417** | pAc5.1-mEGFP-SoYb | EL100 |
| | pAc5.1-mEGFP-**SoYb 1418-1476** | pAc5.1-mEGFP-SoYb | EL101 |
| | pAc5.1-mEGFP-**Vreteno** iso A | *Drosophila* ovarian cDNA | JM59 |
| | pAc5.1-mEGFP-**Yb** iso A | *Drosophila* ovarian cDNA | CB18 |
| **pAc5.1-mEGFP (C-ter)** (FspAI) (EB02) | pAc5.1-**Zucchini***-mEGFP | Sequence specific deletion in pAc5.1-Zucchini-mEGFP | HK171 |
| **pAc5.1-mCherry** (EcoRV) (T7-MJ) | pAc5.1-mCherry-**Daed** iso A | *Drosophila* ovarian cDNA | HK169 |
| | pAc5.1-mCherry-**Gasz** iso A | *Drosophila* ovarian cDNA | XH02 |
| | pAc5.1-mCherry-**Gasz*-Daed** (Gasz lacks the MLS: ∆415-461) | Gibson assembly using pAc5.1-mCherry-Gasz and pAc5.1-mCherry-Daed | HK340 |
| | pAc5.1-mCherry-**Gasz*-Daed MUT** (I69E/Y98E) | Site directed mutagenesis of pAc5.1-mCherry-Gasz 1-414_Daed 1-279 | HK339 |
| | pAc5.1-mCherry-**Gasz*-Daed DEL** (Gasz∆1-236) | Sequence specific deletion in pAc5.1-mCherry-Gasz 1-414_Daed 1-279 | HK338 |
| **pAc5.1-mCherry (C-ter)** (FspAI) (JM65) | pAc5.1-**Zucchini**-mCherry iso A | *Drosophila* ovarian cDNA | EB11 |
| **pAc5.1-PH-mCherry** (FspAI) (HK49) | pAc5.1-PH-mCherry-**Ago3** isoD | *Drosophila* ovarian cDNA | JM138 |
| | pAc5.1-PH-mCherry-**Armitage** iso B | *Drosophila* ovarian cDNA | JM58 |
| | pAc5.1-PH-mCherry-**Aubergine** iso A | pAc5.1-EGFP-Aubergine | RR137 |
| | pAc5.1-PH-mCherry-**BoYb** iso A | *Drosophila* ovarian cDNA | JM82 |
| | pAc5.1-PH-mCherry-**Gasz*-Daed*** (Gasz and Daed lack the MLS: Gasz∆415-461, Daed∆237-279) | Gibson assembly using pAc5.1-mCherry-Gasz and pAc5.1-mCherry-Daed | HK337 |
| | pAc5.1-PH-mCherry-**Hen1** iso A | *Drosophila* ovarian cDNA | HK146 |
| | pAc5.1-PH-mCherry-**Krimper** iso A | *Drosophila* ovarian cDNA | JM141 |
| | pAc5.1-PH-mCherry-**Maelstrom** iso C | *Drosophila* ovarian cDNA | HK147 |
| | pAc5.1-PH-mCherry-**Maelstrom 1-63** | pAc5.1-PH-mCherry-Maelstrom | XH38 |

| | | | |
|---|---|---|---|
| | pAc5.1-PH-mCherry-**Maelstrom 64-333** | pAc5.1-PH-mCherry-Maelstrom | XH39 |
| | pAc5.1-PH-mCherry-**Maelstrom 333-412** | pAc5.1-PH-mCherry-Maelstrom | EL16 |
| | pAc5.1-PH-mCherry-**Maelstrom 334-459** | pAc5.1-PH-mCherry-Maelstrom | XH40 |
| | pAc5.1-PH-mCherry-**Maelstrom 413-459** | pAc5.1-PH-mCherry-Maelstrom | EL17 |
| | pAc5.1-PH-mCherry-**MaelstromΔ1-63** | Sequence specific deletion in pAc5.1-PH-mCherry-Maelstrom | EL13 |
| | pAc5.1-PH-mCherry-**MaelstromΔ64-333** | Sequence specific deletion in pAc5.1-PH-mCherry-Maelstrom | EL14 |
| | pAc5.1-PH-mCherry-**MaelstromΔ333-412** | Sequence specific deletion in pAc5.1-PH-mCherry-Maelstrom | EL24 |
| | pAc5.1-PH-mCherry-**MaelstromΔ334-459** | Sequence specific deletion in pAc5.1-PH-mCherry-Maelstrom | EL12 |
| | pAc5.1-PH-mCherry-**MaelstromΔ413-459** | Sequence specific deletion in pAc5.1-PH-mCherry-Maelstrom | EL25 |
| | pAc5.1-PH-mCherry-**Mael MUT** (V443E/F448E/V450E) | Site directed mutagenesis of pAc5.1- PH-mCherry-Maelstrom | EL115 |
| | pAc5.1-PH-mCherry-**Nibbler** iso A | *Drosophila* ovarian cDNA | HK158 |
| | pAc5.1-PH-mCherry-**Papi** iso C | *Drosophila* ovarian cDNA | JM142 |
| | pAc5.1-PH-mCherry-**Qin** iso B | *Drosophila* ovarian cDNA | JM149 |
| | pAc5.1-PH-mCherry-**SoYb** iso B | *Drosophila* ovarian cDNA | JM86 |
| | pAc5.1-PH-mCherry-**Spindle E** iso A | pAc5.1-EGFP-Spindle-E | RR129 |
| | pAc5.1-PH-mCherry-**Spindle E MUT** (K339E/V342E) | Site directed mutagenesis of pAc5.1-PH-mCherry-Spindle E | DB36 |
| | pAc5.1-PH-mCherry-**Tapas** iso D | pAc5.1-EGFP-Tapas | RR133 |
| | pAc5.1-PH-mCherry-**Tejas** iso A | pAc5.1-EGFP-Tejas | RR117 |
| | pAc5.1-PH-mCherry-**Tejas SIR** (313-355) | pAc5.1-PH-mCherry-Tejas | RR120 |

| | | | |
|---|---|---|---|
| | pAc5.1-PH-mCherry-**Tejas 1-100** | pAc5.1-PH-mCherry-Tejas | RR126 |
| | pAc5.1-PH-mCherry-**Tejas 96-355** | pAc5.1-PH-mCherry-Tejas | RR127 |
| | pAc5.1-PH-mCherry-**Tejas 334-559** | pAc5.1-PH-mCherry-Tejas | EL76 |
| | pAc5.1-PH-mCherry-**Tejas 356-559** | pAc5.1-PH-mCherry-Tejas | RR128 |
| | pAc5.1-PH-mCherry-**Tejas ΔSIR** (Δ313-355) | Sequence specific deletion in pAc5.1-PH-mCherry-Tejas | RR125 |
| | pAc5.1-PH-mCherry-**Tejas Δ1-100** | Sequence specific deletion in pAc5.1-PH-mCherry-Tejas | RR122 |
| | pAc5.1-PH-mCherry-**Tejas Δ96-355** | pAc5.1-PH-mCherry-Tejas | RR123 |
| | pAc5.1-PH-mCherry-**Tejas Δ356-559** | Sequence specific deletion in pAc5.1-PH-mCherry-Tejas | RR118 |
| | pAc5.1-PH-mCherry-**Tejas Δ334-559** | Sequence specific deletion in pAc5.1-PH-mCherry-Tejas | EL84 |
| | pAc5.1-PH-mCherry-**Tejas MUT** (V530E/V532E) | Site directed mutagenesis of pAc5.1-PH-mCherry-Tejas | JM259 |
| | pAc5.1-PH-mCherry-**Tudor** iso A | (Salgania et al. 2024) | HK130 |
| | pAc5.1-PH-mCherry-**Vasa** iso A | pAc5.1-EGFP-Vasa | RR130 |
| | pAc5.1-PH-mCherry-**Vasa open** (K295N) | Site directed mutagenesis of pAc5.1-PH-mCherry-Vasa | RR132 |
| | pAc5.1-PH-mCherry-**Vasa closed** (E400Q) | Site directed mutagenesis of pAc5.1-PH-mCherry-Vasa | RR131 |
| | pAc5.1-PH-mCherry-**Vreteno** iso A | pAc5.1-mEGFP-Vreteno | HK115 |
| | pAc5.1-PH-mCherry-**Vreteno 1-178** | pAc5.1-mEGFP-Vreteno | EL64 |
| | pAc5.1-PH-mCherry-**Vreteno 178-506** | pAc5.1-mEGFP-Vreteno | EL58 |
| | pAc5.1-PH-mCherry-**Vreteno 179-691** | pAc5.1-mEGFP-Vreteno | EL65 |
| | pAc5.1-PH-mCherry-**Vreteno 511-691** | pAc5.1-mEGFP-Vreteno | CB10 |
| | pAc5.1-PH-mCherry-**Vreteno 1-178 RRM_MUT** (Y69E/M74E/Y116E) | Site directed mutagenesis of pAc5.1-PH-mCherry-Vreteno | MD13 |

| | | | |
|---|---|---|---|
| | pAc5.1-PH-mCherry-**Vreteno 179-691 eTud1_MUT** (Y351E/M358E) | Site directed mutagenesis of pAc5.1-PH-mCherry-Vreteno | MD15 |
| | pAc5.1-PH-mCherry-**Vreteno RRM_MUT** (Y69E/M74E/Y116E) | Gibson assembly using pAc5.1-PH-mCherry-Vreteno 1-178 RRM_MUT (Y69E/M74E/Y116E) and pAc5.1-PH-mCherry-Vreteno | DB35 |
| | pAc5.1-PH-mCherry-**Vreteno eTud1_MUT** (Y351E/M358E) | Gibson assembly using pAc5.1-PH-mCherry-Vreteno 179-691 eTud1_MUT (Y351E/M358E) and pAc5.1-PH-mCherry-Vreteno | DB38 |
| | pAc5.1-PH-mCherry-**Vreteno RRM/eTud1_MUT** (Y69E/M74E/Y116E/ Y351E/M358E) | Gibson assembly using pAc5.1-PH-mCherry-Vreteno 1-178 RRM_MUT (Y69E/M74E/Y116E) and pAc5.1-PH-mCherry-Vreteno 179-691 eTud1_MUT (Y351E/M358E) | DB39 |
| | pAc5.1-PH-mCherry-**Yb** iso A | *Drosophila* ovarian cDNA | JM81 |
| **pAc5.1-λN-HA** (EcoRV) (T8-MJ) | pAc5.1-λN-HA-**Tejas** iso A | *Drosophila* ovarian cDNA | I23-MJ |
| | pAc5.1-λN-HA-**Krimper** iso A | *Drosophila* ovarian cDNA | JM168 |
| | pAc5.1-λN-HA-**Vreteno** iso A | pAc5.1-mEGFP-Vreteno | HK133 |
| **pAc5.1A** (EcoRV) (T4-MJ) | pAc5.1-**Armitage** iso B | pAc5.1-mEGFP-Armitage | MD02 |
| | pAc5.1-**Maelstrom** iso C | Deletion of mEGFP sequence from pAc5.1-mEGFP-Maelstrom | MD08 |
| | pAc5.1-**Tejas MUT** (V530E/V532E) | pAc5.1-EGFP-Tejas MUT (V530E/V532E) | HK323 |
| **pGEX-6P-1** (SmaI) | pGEX-6P-1-**Tejas** | *Drosophila* ovarian cDNA | A54-MJ |
| | pGEX-6P-1-**sfGFP**-Tejas (GsfG-Tejas) | Gibson assembly to insert sfGFP sequence in pGEX-6P-1-Tejas | KM-04 |
| | pGEX-6P-1-**sfGFP** (GsfG) | Sequence specific deletion of Tejas from pMJ-GST-sfGFP-Tejas | DH13 |

| | | | |
|---|---|---|---|
| **pMJ-His-MBP** (ScaI) (T46-MJ) | pMJ-His-MBP-**Maelstrom** | *Drosophila* ovarian cDNA | MM2 |
| | His-MBP-**Maelstrom Δ413-459** | Sequence specific deletion in pMJ-His-MBP-Maelstrom | KM01 |
| | His-MBP-**Maelstrom TIR** (413-459) | pMJ-His-MBP-Maelstrom | KM02 |
| | His-MBP-**Mael TIR MUT** (V443E/F448E/V450E) | Site directed mutagenesis of His-MBP-Maelstrom 413-459 (TIR) | EL116 |
